# A Scaffolding Element Rewires Local 3D Chromatin Architecture During Differentiation

**DOI:** 10.1101/2024.05.23.595561

**Authors:** Ivana Jerković, Marco Di Stefano, Hadrien Reboul, Michael F Szalay, Davide Normanno, Giorgio L Papadopoulos, Frederic Bantignies, Giacomo Cavalli

## Abstract

Upon differentiation chromatin rewires to reflect its new cellular identity and function. While it is widely known that this process involves cooperative changes in transcription, chromatin composition and 3D conformation, it is unclear what exactly drives these changes and how they influence one another. Here we used ESC-to-NPC differentiation to study rewiring at a 3 Mb large neuronal *Zfp608* locus. During this process, this large chromatin domain splits in half right at the *Zfp608* promoter, local chromatin gets littered with activating marks, compacts in 3D space and *Zfp608* abounds in transcription. We investigated the *cis* and *trans* elements using capture Hi-C (cHi-C), extensive biophysical modelling, and 3-colour 3D-FISH with technical and analytical breakthroughs and found that transcription abundance modulates the contacts in the region as well as the insulation at the domain split. Furthermore, we found a genetic element we named scaffolding element, with a dual enhancer and architectural function that is essential for chromatin rewiring and loop formation at the NPC stage. The loss of this element disrupts the formation of all local NPC-loops irrespective if they are anchored in this element or not, highlighting the hierarchical relationship between elements that act as loop anchors. Furthermore, we uncovered that the scaffolding function, although driven by multiple mechanisms, can form loops independent of loop-extrusion and that other molecular attractions were necessary to form NPC-specific contacts in the region. Together, these results demonstrate that a hierarchy of genetic elements in *cis* allows successful rewiring during differentiation and that multiple *trans* acting elements contribute to make this rewiring efficient.

## 2. Introduction

High cellular plasticity is a hallmark of pluripotent cells allowing them to respond to developmental and differentiation cues. The molecular basis of this response is multifaceted and occurs concomitantly at the epigenomic, transcriptomic and chromatin conformation levels^1–4^. In particular, chromatin conformation changes during cell differentiation processes are well documented and encompass compartment switching, domain restructuring, and enhancer-promoter (E-P) contact rewiring^2–7^. Local contacts rewiring (under 1 Mb) is often associated with specific gene-expression changes, highlighting the importance of chromatin contacts in regulating transcription^3^. However, even though many studies focused on the causal relationship between chromatin contacts and transcriptional response, they gave conflicting results indicating a context-dependent relationship^8–19^. In addition, most of the contacts are not functionally annotated, hence their potential contribution to the transcriptional response is not entirely known. Aside from activating and repressive loops, several recent studies described different types of chromatin contacts (tethering elements and meta-loops) that flank *cis*-regulatory elements and might modulate the timing of E-P interactions and the robustness of transcriptional regulation^7,20–22^. Nevertheless, little is known about the mechanisms regulating specific 3D chromatin contacts, in particular in *cis,* leaving several unresolved questions about the hierarchical nature of these contacts, their importance for transcriptional regulation in steady state and differentiation^7,20,22^.

To study the mechanism and dynamics of these changes, we used an *in vitro* neuronal progenitor differentiation system which closely resembles *in vivo* neuronal differentiation^3,5–7^. In this system, the most striking chromatin reorganization happens at neuronal progenitor cell (NPC) specific genes, like *Zfp608*^3^. The *Zfp608* gene function is relatively elusive. It is expressed in several cell-types like thymocytes where it represses *Rag1* and *Rag2* and thereby suppresses T-cell maturation in the developing fetus^23,24^. In neuronal tissues, this gene has been identified as a potential biomarker for the Cornelia de Lange Syndrome (CdLS), where it works antagonistically with its paralogue *Zfp609* and its partner *Nipbl*^25,26^. Here, we focused on the *Zfp608* locus to study chromatin conformation rewiring during the mouse embryonic stem cell (ESC)−to−NPC differentiation and its connection to transcription. By combining molecular biology, biophysical modelling, and imaging with novel analyses in wildtype (wt) and seven mutant cell lines in ESC and NPC states, we demonstrate that transcription directly contributes to the NPC-specific rewiring via 3D contact insulation and loop stabilization. Furthermore, we discover a regulatory region with a dual functional and architectural role, that is essential for contact rewiring and involves both CTCF-dependent insulation and loop-extrusion as well as CTCF-independent looping interactions.

## 3. Results

### NPC differentiation rewires chromatin composition, transcription, and local chromatin contacts

To investigate cell-type specific chromatin and transcriptional rewiring at the *Zfp608* locus we differentiated wt ESCs into NPCs and characterized their chromatin composition (ChIP-seq of H3K27Ac, H3K4me3, H3K36me3, Pol2Pol2, PAX6, CTCF and Rad21), transcriptional status (RNA-seq), and local 3D chromatin conformation at the *Zfp608* locus (cHi-C) in biological replicates (**Fig. 1a, Extended Data Fig. 1a, 2a).** NPC differentiation is accompanied by active histone mark deposition at the locus, *Zfp608* transcriptional activation, and by 3D chromatin contact rewiring (**Fig. 1a, 1b, Extended Data Fig. 2b, Supplementary Table 1, and 2)**^3,5^. *Zfp608* is located in a large chromatin domain (∼3 Mb) devoid of other genes that splits into two upon NPC differentiation due to a significant increase in insulation at the *Zfp608* promoter (**Fig. 1a, 1b, Supplementary Table 1, and 2**). This boundary formation is concomitant with the emergence of multiple, specific chromatin contacts (loops) that appear mainly upstream to the gene promoter **(Fig. 1c, and 1d).** These NPC-specific contacts include promoter anchored loops (putative E-P loops) and contacts among elements covered by the H3K27Ac mark and bound by the PAX6 transcription factor (putative E-E loops). Notably, apart from the promoter region and one putative enhancer region, which we named element A, we did not observe CTCF/RAD21 binding in any other loop anchor (**Fig. 1b, 1c, Extended Data Fig. 3a and 3b**). To examine the main NPC-specific putative E-P contacts and their importance to the establishment of the NPC-specific domain we selected two promoter-distal loop anchors based on their differential TF occupancy and histone mark depositions. One loop anchor was a putative enhancer, element A and the other a loop anchor largely devoid of any marks or TFs, element B **(Fig. 1a, and** Extended Data Fig. 3b). According to the cHi-C maps, both of these two elements contact the *Zfp608* promoter and other putative enhancers over long distances (560 kb and 826 kb to the promoter), but are occupied by a distinct set of TFs and chromatin marks. In NPCs, element A harboured H3K27Ac, PAX6, Pol2, CTCF and RAD21 bindings whereas element B, only exhibited weak PAX6 occupancy and very weak H3K27Ac signal (**Fig. 1b, and** Extended Data Fig 3b). Next, we tested the enhancer activity potential of these two elements by luciferase assay in three different cell lines (ESC wt, NPC wt, and ESC cell line ectopically expressing *Pax6* (ESC Pax6+)) and using two different promoters (SV40 or cognate *Zfp608* promoter) (**Fig. 1e and Extended Data Fig. 3c-f)**. We found that, in every cell line and using either promoter, only element A had enhancer activity **(Fig. 1e and Extended Data Fig. 3c-f).** Altogether, these data demonstrate that *Zfp608* change in transcriptional activity is accompanied by chromatin conformation rewiring including changes at multiple putative regulatory elements, one of which, element A, behaves as an enhancer that establishes multiple E-P and P-P long-range chromatin contacts.

**Figure 1.**
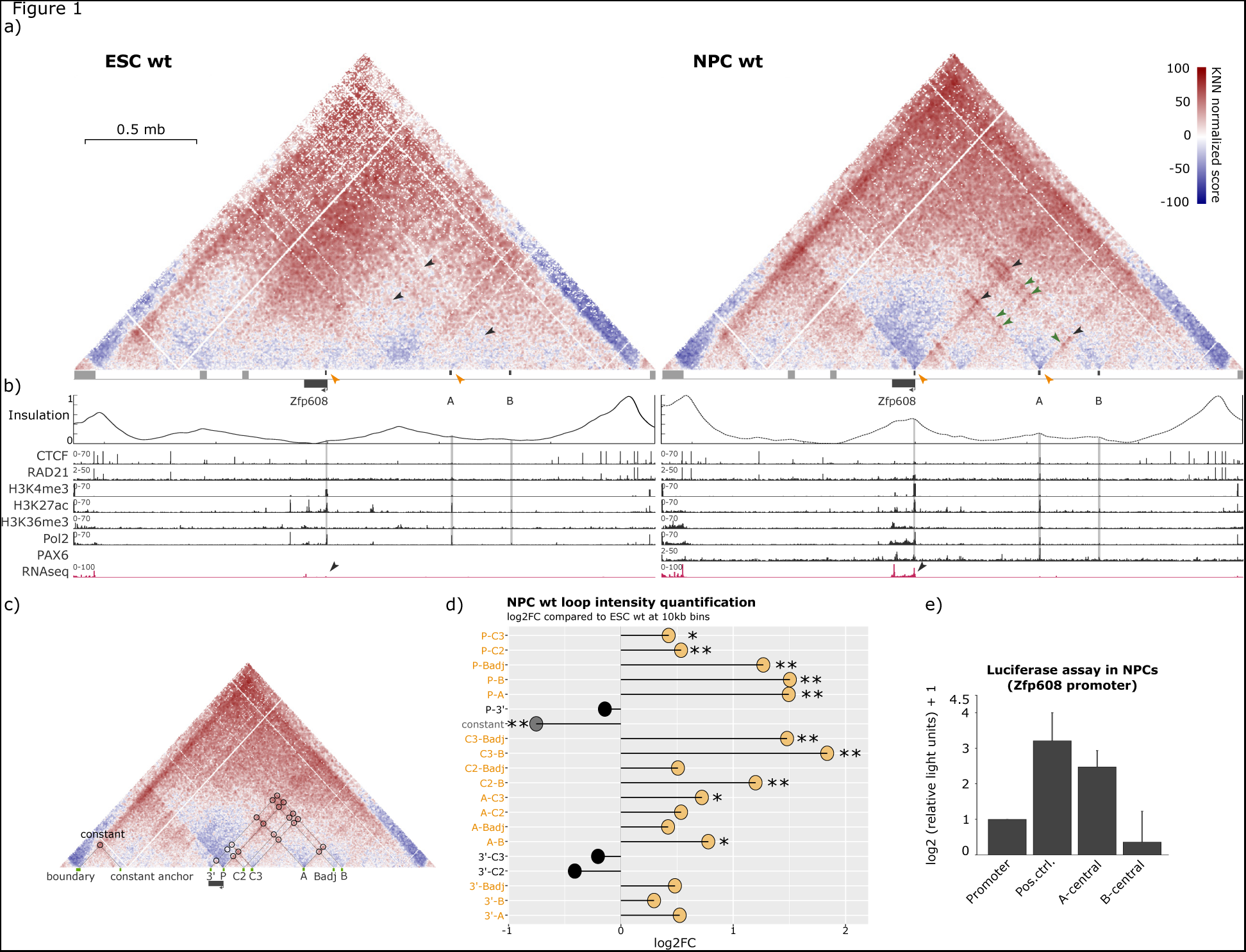
Experimental Capture Hi-C (cHi-C) maps at the *Zfp608* locus in ESC and NPC wt cells. **a)** Score (observed/expected, see **Methods**) maps from cHi-C experiments in ESC and NPC wt cells at 5 kb resolution. Three NPC-specific contacts of interest are annotated with black arrows: promoter-A (P-A), promoter-B (P-B), and (A-B). Green arrows point to other NPC-specific putative enhancer-enhancer (E-E) contacts, and orange arrows annotate two main NPC-specific insulation points at the *Zfp608* promoter and element A. **b)** From top to bottom: insulation score (IS), ChIP-seq (CTCF, RAD21, H3K4me3, H3K27Ac, H3K36me3, Pol2 and PAX6), and RNA-seq profiles at the capture region for ESC and NPC states. Black arrows point to the RNA-seq signal at the *Zfp608* promoter. **c)** Circles on an NPC wt cHi-C map show the main NPC-specific putative E-P and E-E contacts. Below the map, the anchors of each contact (loop) are named consistently with the annotation in panel d (*i.e.*, P-A is a loop with one anchor in P and the other in A). **d)** log2-fold-change (log2FC) of the loop intensity in NPC wt over the ESC wt condition. Loops with increased strength in NPCs are shown in orange. * indicates 0.01<*p*<0.05, and ** indicates *p*<0.01, where the p-values of the log2FC were calculated using a Wald statistical test using DEseq2 (**Methods**). **e)** Luciferase assay in NPC cells with: a cognate *Zfp608* promoter and no enhancer (Promoter), a cognate *Zfp608* promoter with a positive control enhancer (SV40 enhancer) (Pos.ctrl.), a cognate *Zfp608* promoter with a central part of element A (A-central) and a cognate *Zfp608* promoter with a central part of element B (B-central).

### Transcription attenuation weakens insulation and contacts at the *Zfp608* promoter

To understand the interplay of 3D genome organization and transcription activity at the *Zfp608* locus we tested how perturbing the *Zfp608* transcription start site (TSS) and transcription strength impacts the local chromatin structure. To this end, we engineered two mutant cell lines: a proximal promoter deletion (Δprom), and a 3xSV40 polyA insertion (polyA) immediately downstream of the main TSS (**Fig. 2a and Methods).** As for the wt case, we profiled the local 3D chromatin conformation (cHi-C), transcriptional status (RNA-seq) and chromatin composition (ChIP-seq of H3K27Ac, H3K4me3, H3K36me3, RNA Pol2 (Pol2), PAX6, CTCF and Rad21) in both mutants in ESC and NPC state. The two mutants differ substantially from one another in design. In the Δprom mutant, a proximal part of the promoter and first two exons are deleted (∼4.3 kb), which directly impacts TF binding and histone marks at the promoter (**Fig. 2a, and Extended Data Fig. 54a**). In the polyA mutant, the SV40 signal is inserted 78 bp downstream to the main TSS leaving the TFs binding, histone marks, and the TSS position intact (**Extended Data Fig. 4a**). In this way, we could assess and uncouple the effect of the transcription from the effect of the chromatin factor binding at the promoter. We found no changes at the *Zfp608* locus in terms of transcription, chromatin conformation, TF binding or histone marks in any ESC mutant line compared to wt (**Extended Data Fig. 1b, 1c, 1d, 2a, 4c, Supplementary Table 1, and 2**). Interestingly, in the NPC state, Δprom and polyA mutants affect transcription differently. In the Δprom mutant, transcription starts upstream of the deleted TSS and successfully transcribes the rest of the non-deleted gene part. This causes overall mild and not significant transcriptional changes (**Fig. 2a, 2b, 2c, Extended Data Fig. 2c, 4a, 4b**, and **4c)**. In contrast, polyA significantly reduces transcription (to ∼25-50% (qPCR) and ∼50% (RNA-seq)) of the NPC wt level) over the entire gene body (**Fig. 2a, 2b, 2c, Extended Data Fig. 4a, 4b**, and **4c**). Notably, none of the changes in transcription were mirrored by changes in TF occupancy or in histone modifications at the *Zfp608* locus (**Fig. 2a, and 2b**).

**Figure 2.**
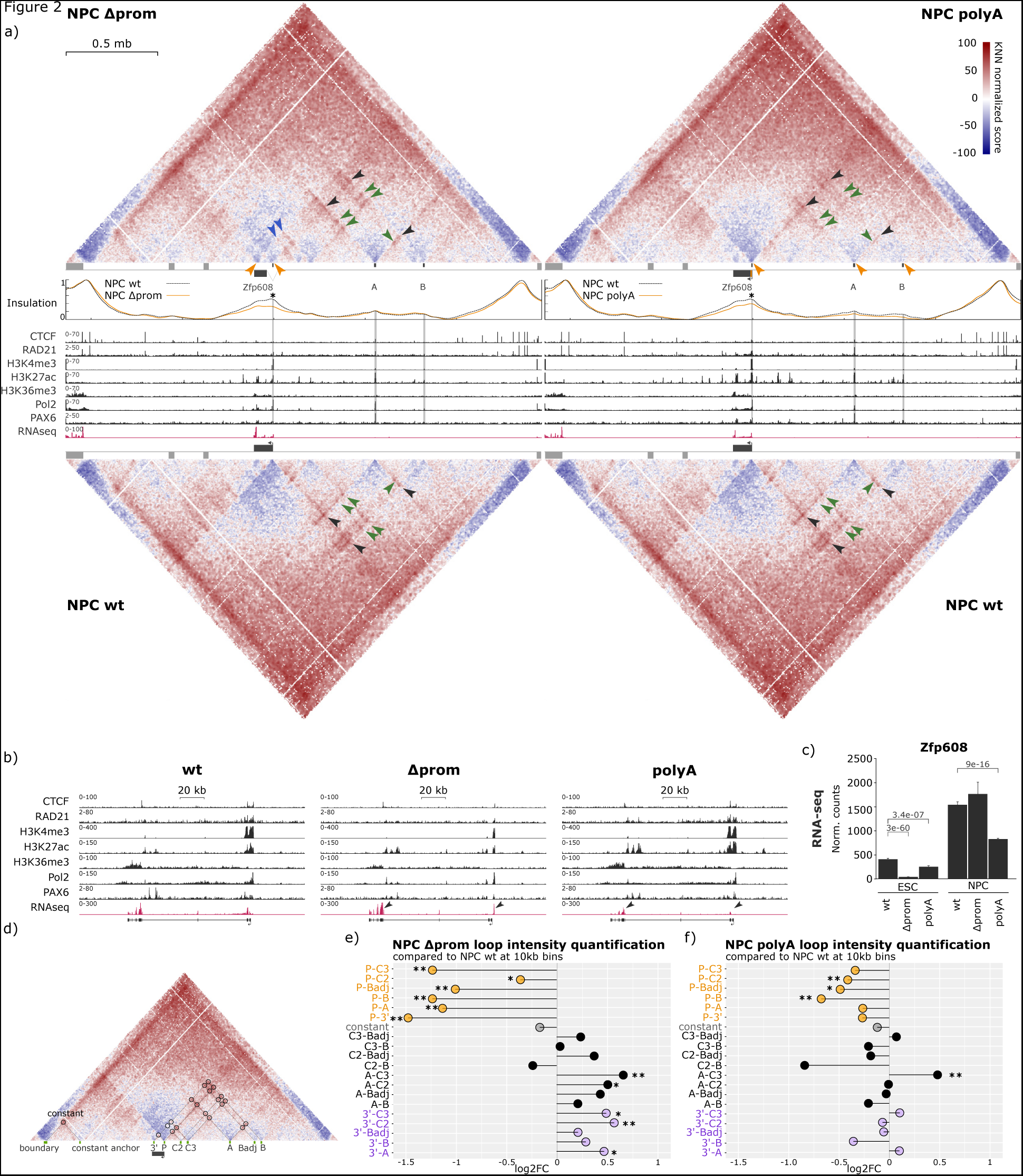
Impact of transcription on local chromatin conformation. a) NPC cHi-C score maps (observed/expected) at 5 kb resolution for the wt, partial promoter deletion (Δprom, 4.3 kb deletion), and polyA insertion (3x SV40 polyA inserted 78 bp under the main TSS) conditions (**Methods**). Three NPC-specific contacts (P-A, P-B, and A-B) are annotated with black arrows. Green arrows point to NPC-specific putative E-E contacts. Blue arrows indicate ectopic contacts and orange arrows indicate sites of insulation changes in NPC Δprom and polyA in comparison to the NPC wt. Below cHi-C maps, the top panel depicts the experimentally quantified insulation score at the region. The asterisk indicates a significantly changed insulation score between mutant and wt condition calculated by Welch Two Sample t-test: Δprom pval=1.26E-09, and polyA pval=1.3E-04 (**Methods**). The panels below show ChIP-seq (CTCF, RAD21, H3K4me3, H3K27Ac, H3K36me3, Pol2, and PAX6), and RNA-seq data. **b)** Zoom-in on the *Zfp608* gene with the ChIP-seq and RNA-seq data as in a). Black arrows point to the RNA-seq profile in the 3’ and 5’ ends of the gene in the two mutant conditions. In Δprom, they indicate a new transcription start site, the missing first two exons, and a slightly more abundant signal at the end of the transcript. In polyA, the arrows indicate a relatively even reduction in transcript abundance. **c)** Normalized RNA-seq counts of ESC and NPC wt, Δprom, and polyA cell lines and their p-adjusted (p_adj_) values from Deseq2 analysis. Each p_adj_ value (Wald test) was calculated by pairwise comparison of each mutant to the wt of the corresponding cell line (*i.e.*, ESC wt to ESC polyA) (**Methods** and **Supplementary Table 2**). **D)** A schematic of an NPC wt map with NPC-specific putative E-P and E-E contacts. Below the map, the anchors of each contact (loop) are named consistently with the annotation in panels e-f (i.e., P-A is a loop with one anchor in P and the other in A). **e)** and **f)** log2FC of the loop intensity in each of the mutants over the NPC wt condition (**Methods**). Loops that have one anchor at the promoter (P) are annotated in orange and (ectopic) loops that cross the promoter (high point of insulation) are annotated in purple. * indicates loops with 0.01<*p*<0.05, and ** indicates *p*<0.01, where the *p*-values of the log2FC were calculated with a Wald statistical test using DEseq2 (**Methods**).

Next, we analysed the cHi-C maps, in particular the local chromatin contacts and insulation (**Fig. 2a**). We found that, despite different effects on transcription, both mutants significantly decreased insulation at the promoter and at the 3’ end of the *Zfp608* gene (**Fig. 2a, orange arrows and asterisk).** We notice that insulation was also slightly reduced at elements A and B in the polyA mutant (**Fig. 2a, orange arrows**). In addition, we found ectopic contacts (newly present in the mutant) crossing the *Zfp608* promoter (a high point of insulation in wt) in Δprom but not in polyA, likely due to lower insulation in Δprom mutant (**Fig. 2a (blue arrows), Fig. 2a, 2e, and 2f, purple labelled contacts**). Furthermore, we found that the promoter-anchored contacts significantly weakened in both mutants (**Fig. 2e, and 2f**, **orange labelled contacts**). This effect was stronger in the Δprom mutant, consistent with the fact that the deletion overlaps with the loop anchor site (**Fig. 2b, 2e, and 2f**). Interestingly, promoter-unrelated (putative E-E) contacts were affected differently in the two mutants as they were reinforced in the Δprom but not in the polyA mutant (**Fig. 2d, 2e, and 2f, purple and black labelled contacts**).

These findings show that the partial loss of histone marks and TF binding (Δprom) at the promoter as well as the attenuated transcription strength (polyA) directly affect the establishment of the NPC-specific domain. In Δprom, contacts and insulation might be attenuated due to the missing transcription factor binding and altered transcription initiation site, while in polyA this is exclusively due to the decrease in transcription which potentially destabilizes and weakens chromatin loops^17^. Altogether, these findings highlight a rather dynamic relationship between structure and transcription which influence each other.

### Chromatin contacts are hierarchically organized

Next, we characterized the role of the NPC-specific putative E-P loops and their contribution to rewiring. We focused on the elements A and B and engineered three different cell lines where we deleted element A (ΔA), element B (ΔB), or both (ΔAB). We, then, profiled them by cHi-C, RNA-seq and ChIP-seq in ESCs and NPCs (**Fig. 3a, 3b, Extended Data Fig. 1e-g, 5a, and 5b**). In the ESC state, we found no significant changes in *Zfp608* expression, in the ChIP-seq profiles or in the cHi-C profiles in any of the mutant cell lines (**Extended Data Fig. 1e-1g, 4f, 5e, Supplementary Table 1, and 2**). Moreover, even though element A functions as an enhancer in transfection assays, neither deletion significantly perturbed *Zfp608* expression (**Fig. 3c, Extended Data Fig. 2d, and 4d-f**). Since the *Zfp608* region is littered with putative enhancers, which never completely lose interaction with the promoter, it is likely that there is some functional redundancy between these elements and that individual enhancer deletions are insufficient to affect transcription.

**Figure 3.**
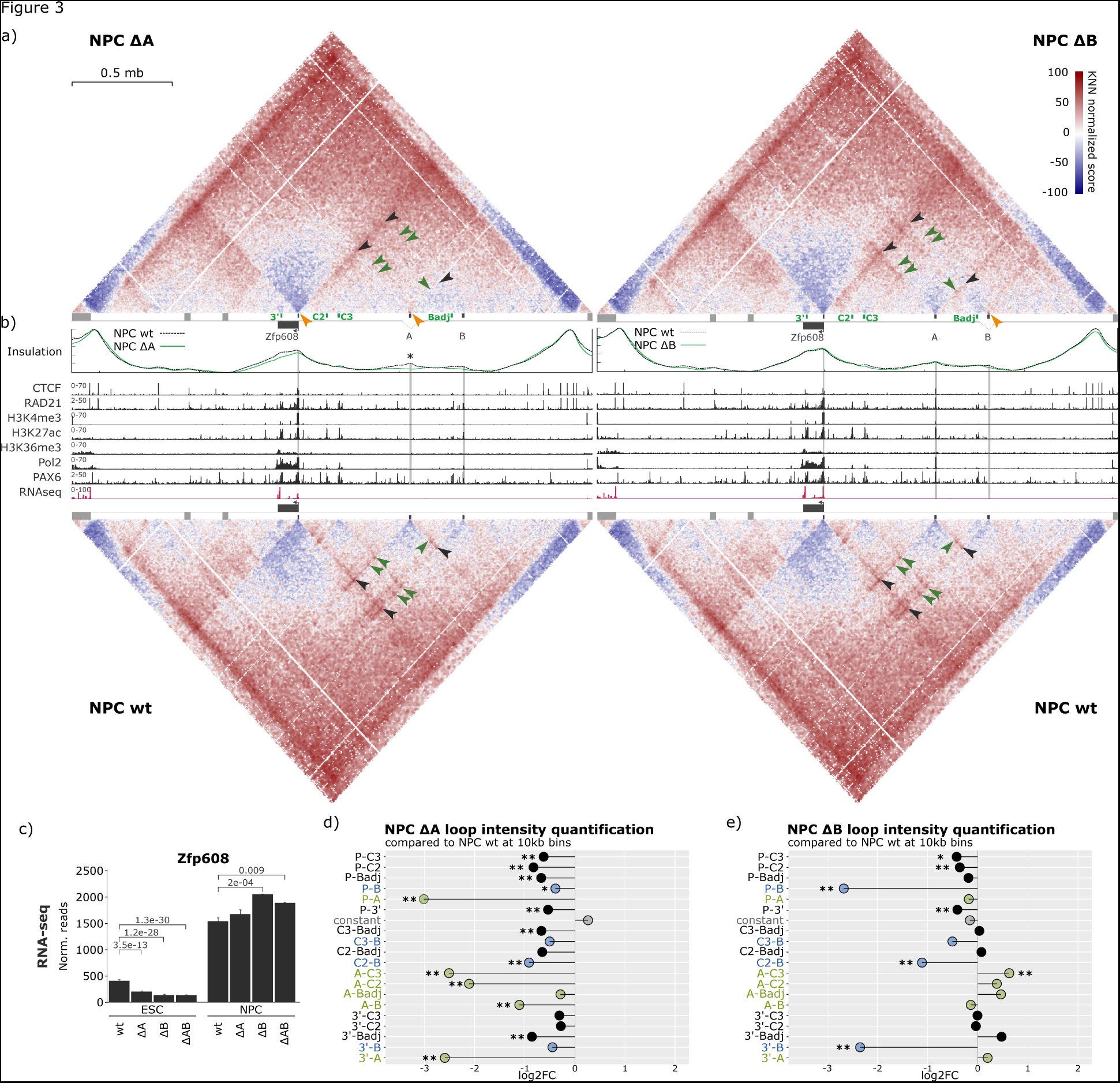
Impact of distinct NPC-specific loops on local chromatin conformation. **a)** NPC cHi-C score maps (observed/expected) of the wt, element A deletion (ΔA, 8.2 kb deletion), and element B deletion (ΔB; 7.4 kb deletions) are shown at 5 kb resolution. Three NPC-specific contacts are annotated with black arrows (P-A, P-B, and A-B) and green arrows indicate the visible retention of some putative E-E contact in ΔB but not ΔA mutant. **b)** The top panel depicts experimentally quantified insulation score (IS) profiles for ΔA or ΔB (light green), and wt (black dashed line) conditions. The asterisk indicates sites of significantly changed IS between mutant and the wt condition calculated by Welch Two Sample t-test; ΔA condition at enhancer A - pval=3.1E-13. The panel below shows ChIP-seq (CTCF, RAD21, H3K4me3, H3K27Ac, H3K36me3, Pol2, and PAX6), and RNA-seq data. **c)** Normalized RNA-seq counts of ESC and NPC states of wt, ΔA, and ΔB, and ΔAB cell lines and their p_adj_ values from Deseq2. Each p_adj_ was calculated with a Wald test by pairwise comparison of individual mutants to the wt of the corresponding cell line (*i.e.*, ESC wt to ESC ΔA). **d)** and **e)** log2FC of the loop intensity in each of the mutants over the NPC wt condition (**Methods**). All loops that have one anchor in A are annotated in green and all loops that have one anchor in B are annotated in blue. * indicates loops with 0.01<*p*<0.05, and ** indicate *p*<0.01, where the *p*-values of the log2FC were calculated from a Wald statistical test (**Methods**).

Conversely, the 3D genome organization was perturbed to varying degree in all three mutants. Strikingly, the ΔA mutant (∼8.2 kb deletion) significantly affected not only the loops anchored in this element but also all other enhancer A-independent loops as well as most promoter-anchored loops at the *Zfp608* locus (**Fig. 3a, and 3d**). In addition, this deletion significantly changed the insulation at element A but not at the *Zfp608* promoter (**Fig. 3b, asterisk above the insulation**). The double deletion (ΔAB) affected local architecture similarly to the ΔA condition (**Extended Data Fig. 5a, and 5d**). In contrast, the element B deletion (∼7.4 kb deletion) caused a significant attenuation of the element B-anchored contacts but had no significant impact on the insulation nor on the element B-independent contacts in the region (**Fig. 3a, 3b, and 3e).** These findings suggest that element A may have a local master architectural role, bringing the other elements at the locus in 3D proximity. This could happen either because A is a contact hub and directly interacts with multiple elements simultaneously, or because A is a scaffolding element, where a single frequent contact, possibly with the *Zfp608* promoter, increases the likelihood of further chromatin contacts by bringing other elements in 3D proximity.

### Element A scaffolds the region upon differentiation

To test these two hypotheses, we performed 3-color 3D-FISH in ESC wt, NPC wt, and NPC ΔA mutant to quantify simultaneously in single-cells the distances between promoter (P), element A (A), and element B (B). For this, we developed small, highly sensitive oligopaint probes (**see Methods**) to be able to balance the signal strength and the sensitivity of the experiment. Specifically, we designed three 10 kb probes covering P, A, and B and a 15 kb probe for the element A in ΔA mutant condition (two 7.5 kb probes flanking each side of the deletion) (**Fig. 4a, and Methods**). Then, we analysed pairwise distances as well as the triplet distances among P, A and B elements (**Fig. 4b and 4c**).

**Figure 4.**
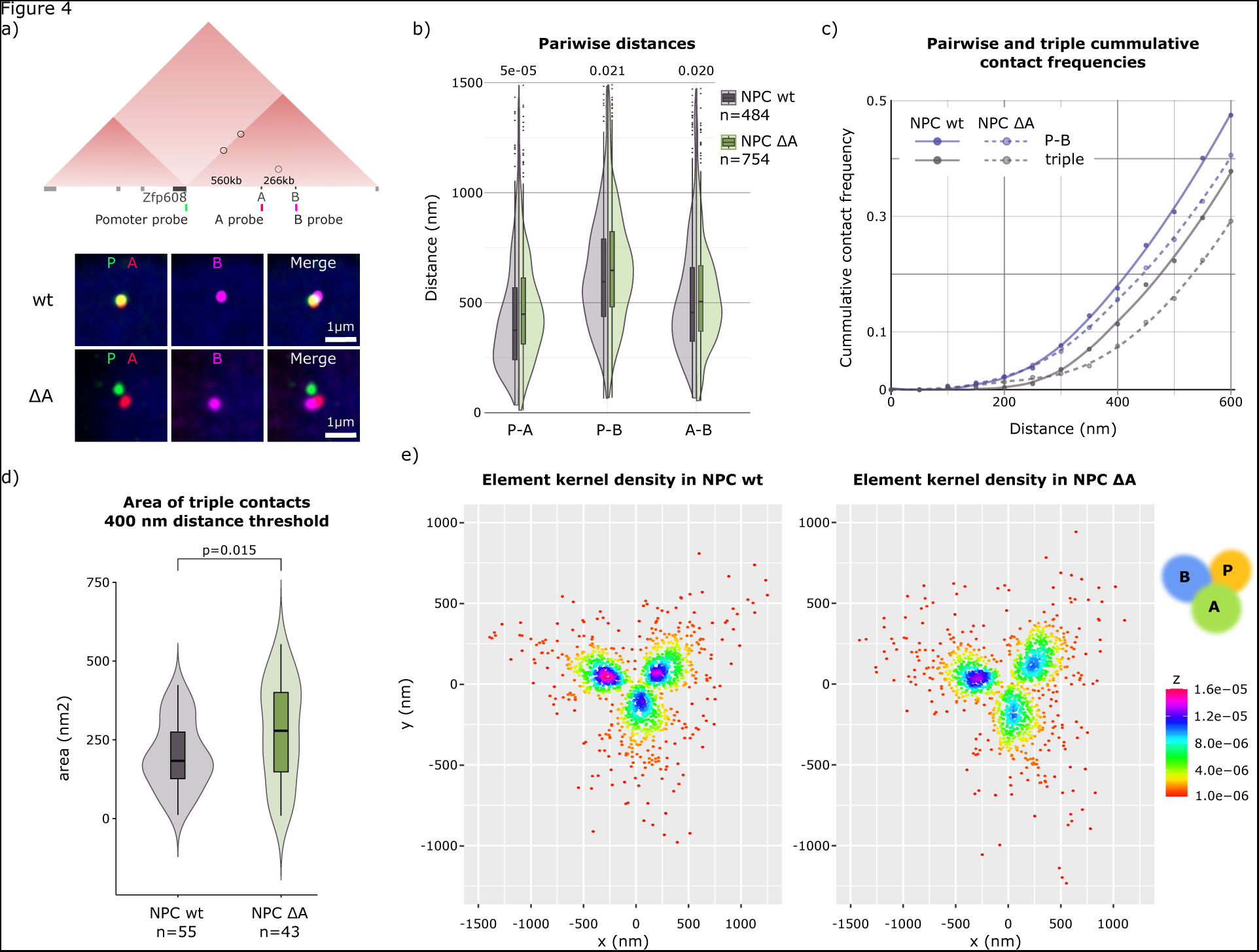
3-color 3D-FISH in NPC wt and NPC ΔA conditions. **a)** Schematic representation of the 3-colour 3D-FISH probe positions (promoter (P), A and B) and their mutual genomic distances and representative images for the two conditions. **b)** Pairwise distance distributions between P-A, P-B, and A-B in NPC wt and NPC ΔA **(Methods)**. p-values were calculated using the Mann-Whitney statistical test comparing wt with ΔA. **c)** Pairwise P-B (blue) and triple (P-A-B, grey) cumulative contact frequencies as a function of the distance threshold for NPC wt (solid line) and NPC ΔA (dashed line) cells **(Methods)**. Triple-contacts are determined if all of the three pairwise distances are lower than a certain threshold (i.e., if P-A, A-B, and P-B distances are all under 400 nm, then this is considered a triple-contact at 400 nm). Each point shows where the contact frequency has been calculated and the line is a regression line to help to follow the tendency. **d)** Area of the triangle formed by the three imaged points (P, A, and B) at 400 nm distance threshold **(Methods** and **Extended Data** Fig. 6c), p-value was calculated using the Mann-Whitney statistical test. **e)** Kernel density plots of the P, A, and B positions in NPC wt and ΔA. The kernel density at each pixel represents the density of the elements and is shown with a colour code (**Methods** and **Extended Data** Figure 6d**, 6e, and 6f)**. Each plot can be read as a topographical map where the highest density values in purple and pink represent the highest peaks on the map.

We found that in NPCs all three pairwise distances (P-A, P-B, and A-B) are significantly shorter compared to the ESC wt condition indicating that the locus is compacting upon differentiation (**Extended Data Fig. 6a)**. In addition, the closest distance measured was P-A, despite A and B being twice closer in genomic distance (A-P = 560 kb and A-B = 266 kb) (**Extended Data Fig. 6a**). Interestingly, this effect is also visible in ESC wt cells. To investigate how strong is this tendency in the two conditions, we calculated the ratios of median A-B and P-A distances (med(A-B)/med(P-A) in wt ESC and NPC and found that the ratio is larger in NPCs (1.136284) than in ESCs (1.047649) indicating that, beyond a putative effect of global chromatin condensation, the contacts between P and A elements become particularly strong in NPCs (**Extended Data Fig. 6a**).

We, therefore focused on the analysis of the effect of the ΔA mutation in the NPC state. We found that upon deletion of enhancer A, all three pairwise distances were significantly increased, with P-A being most strongly affected, indicating that all A-anchored contacts aren’t affected equally but also that the element A facilitates the A-independent P-B contact (**Fig. 4b**). By varying the distance threshold to detect percentage of contacts between pairs of loci, we found that, in NPC wt, P and B are rarely closer than 200 nm (in <2.3% of cells), 300 nm (7.6%) or even 400 nm (17.6%). Conversely, P and A are found much more often closer than 200 nm (15.9%), 300 nm (35.3%) or 400 nm (51.9%), indicating that P and A interact either more frequently or stably (**Fig. 4c and Extended Data Fig. 6b).**

Next, we quantified how often we find all the three loci in the vicinity of one another (triplet) at varying distances (**Fig. 4c**). The incidence of triple contacts in NPC wt is 0.4% under 200 nm, 3.5% under 300 nm and 11.4% under 400 nm. This implies that 45.9% of the cells that present a P-B distance under 300 nm are also forming a triplet and this percentage increases to 64.7% for distances under 400 nm, respectively, strongly suggesting that the contact between element B and the promoter is facilitated by A to promoter proximity (**Fig. 4c and Extended Data Fig. 6b**). Consistent with this data, the proportion of triplets decreased in ΔA mutant. Triplet frequencies start reducing between the wt and ΔA from the 300 nm distance threshold onward. The proportion of cells with triplets under 300 nm is reduced from 3.5% in wt to 2.8% in ΔA (p-val. = 0.6603, two proportions z test), under the 400 nm from 11.4% in wt to 7.6% in ΔA (p-val. = 0.05659, two proportions z test) and under 450 nm from 18.2% in wt to 11.7% in ΔA (p-val. = 0.00604, two proportions z test). This trend continues with the same margin of difference up until the of 650-700 nm distance when it starts to decrease (**Fig 4c, and Extended Data Fig. 6b**). We note that the reduction of triplets in ΔA mutant reflects changes in P-A and A-B distances, but also in P-B distances. Therefore, the deletion of element A leads to a significant increase in all the pairwise distances P-A, A-B, and P-B, resulting in fewer triplets but not in their complete loss which one would expect if element A behaved as an obligatory contact hub.

To further understand the relationship between the P, A, and B elements, we measured the area of the triangles made by the three elements and compared them between the wt and ΔA. We found that for all distances, the triangle area distribution is significantly higher in ΔA compared to the wt (**Extended Data Fig. 6c**). Even using the 400 nm threshold for triplet identification, we measured a significantly greater surface area in ΔA, showing that in ΔA not only the number of triplets decreases, but even the remaining triplets drift further apart (**Fig. 4d**).

Finally, we set out to investigate how constricted in 3D space are the three loci in NPC wt and the NPC ΔA condition. For this, we used a novel analytical approach where we optimally superimposed the three points on a single plane while preserving their distances. This was done by taking three loci from single cells and *in silico* connecting them to make triangles. The centres of mass of triangles were then stacked on top of one another while maintaining the topological position of the three loci (top left-B, top right-P, and bottom centre-A) (**Extended Data Fig 6d**). Next, we calculated a 2D kernel density of the elements’ positions. Briefly, the same number of occurrences for each element was taken and their density was investigated over the equally sized 2D plane. The density (number of occurrences) of the element is shown as a set of values of each pixel on a 2D map (**Fig. 4e, Extended Data Fig. 6e, and 6f).** We found that, upon the deletion of element A, the chromatin immediately adjacent to A occupies a significantly larger area, indicating that it is less restricted in space (**Fig. 4e, Extended Data Fig. 6f)**. Intriguingly, the same impact is seen on promoter and element B showing that the mutual displacement of the three elements is restricted to a narrower region in NPC wt than in the NPC ΔA (**Fig. 4e**).

Altogether, these experiments suggest that that there is a hierarchical relationship between contacts in this region. Element A acts as an architectural scaffold which, while contacting the promoter, also constrains the movements of other chromatin regions thereby favouring contacts that would otherwise happen less frequently. This constrained dynamic contributes to the efficient chromatin rewiring during differentiation.

### The scaffolding element function mainly relies on loop-extrusion independent mechanisms

Given the scaffolding function of element A, next, we sought to investigate the mechanism driving this behaviour. The inspection of chromatin tracks revealed that element A was the only putative enhancer in this region that displayed the presence of two loop-extrusion proteins, CTCF and RAD21 (cohesin) (**Extended Data Fig. 3b**)^27,28^. To perturb loop-extrusion at A, we engineered a new mutant cell line where we deleted a 22 bp CTCF motif (ΔCTCF_A_) aiming to abolish the CTCF binding and RAD21 accumulation without perturbing the binding of any other factor (**Fig. 5a, and 5b**)^27,28^. Characterization of the ΔCTCF_A_ mutant by cHi-C and ChIP-seq in ESC state revealed no overall architectural, or TF occupancy changes (**Extended Data Fig. 1h**). Conversely, in the NPCs we observed a total loss of CTCF and RAD21 occupancy while the Pol2, H3K27Ac, and PAX6 binding was maintained, albeit to a slightly lower level than in wt with very mild overall transcriptional effect (**Fig. 5a, 5b, 5c**, and **Extended Data Fig. 4e**).

**Figure 5.**
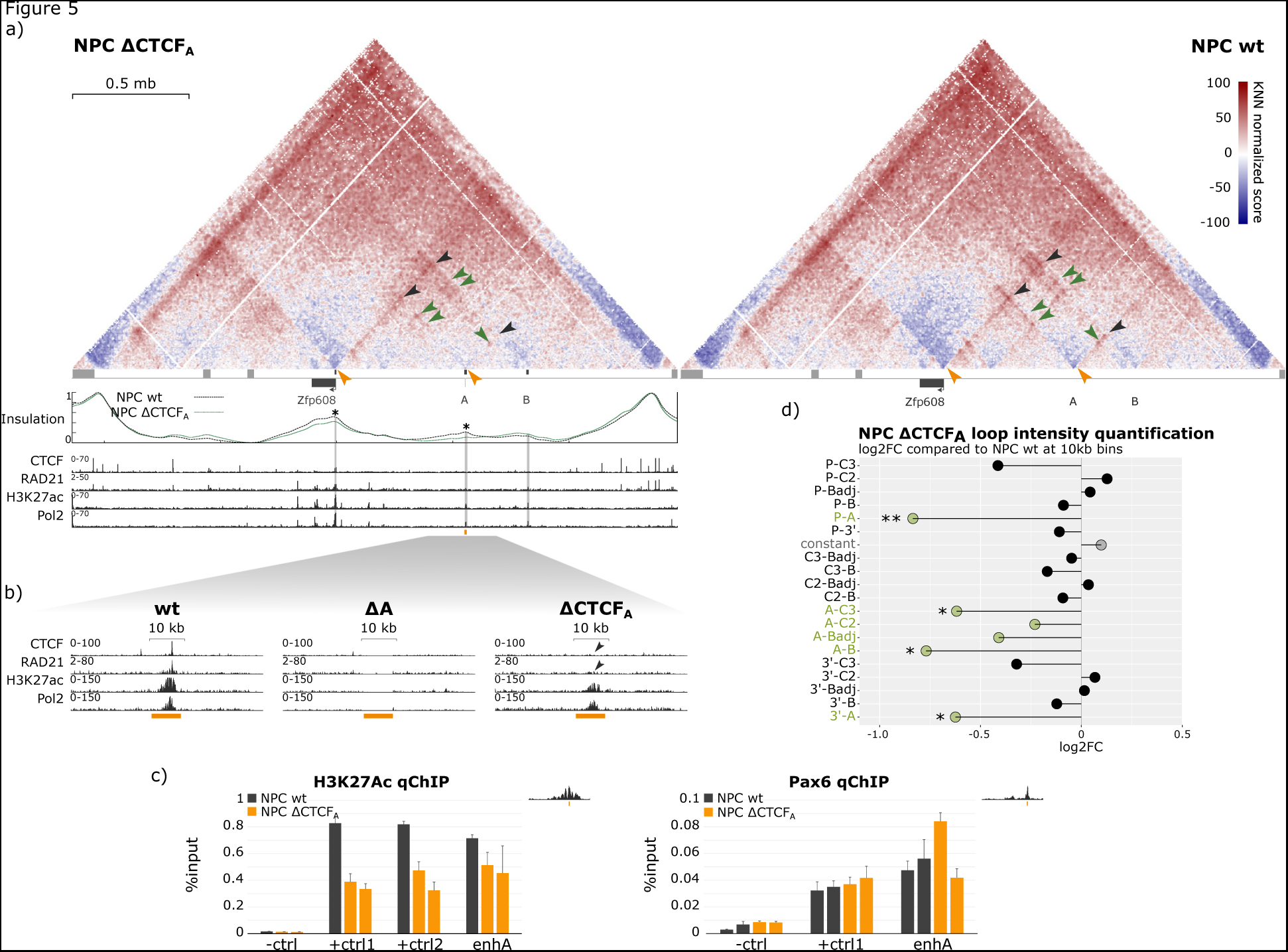
Mechanistic investigation of the scaffolding element A. **a)** NPC cHi-C score maps of the CTCF motif deletion at the element A (ΔCTCF_A_, 22 bp deletion) at 5 kb resolution. Three NPC-specific contacts are pointed to with black arrows (P-A, P-B, and A-B). Green arrows indicate NPC-specific putative E-E contacts that are still present in ΔCTCF_A_ mutant. Orange arrows point to visible changes in insulation pattern between ΔCTCF_A_ and wt. Under the cHi-C map, the top panel depicts the experimentally quantified insulation score (IS) at the capture region. The asterisk indicates significant changes in IS in ΔCTCF_A_ compared to wt where the p-value was calculated by Welch Two Sample t-test (pval=0.002 at promoter, pval=2.6E-10 at A). The panel below shows ChIP-seq (CTCF, RAD21, H3K27Ac, and Pol2) data for ΔCTCF_A_. **b)** Zoom-in at the element A in wt, ΔA, and ΔCTCF_A_. The orange bar indicates the deleted region in the ΔA mutant. Black arrows indicate the loss of CTCF and RAD21 binding in the ΔCTCF_A_ mutant. **c)** H3K27Ac and PAX6 qChIP from NPC wt and ΔCTCF_A_ conditions. -ctrl1 is negative control (a region with no binding of PAX6 or H3K27Ac in their respective ChIPs). +ctrl1 and 2 are positive controls (regions with known binding of PAX6 or H3K27Acin their respective ChIPs). enhA marks the primers at the element A at the peak of expected binding (see the PCR product position (orange) at the NPC wt ChIP peak in the top right corner). **d)** log2FC of the loop intensity in ΔCTCF_A_ mutant in comparison to the NPC wt condition (**Methods**). In green, all loops that have one anchor in A are annotated. * indicates loops with 0.01<*p*<0.05, and ** indicate *p*<0.01, where the *p*-values of the log2FC were obtained using a Wald statistical test in DEseq2 (**Methods**).

The cHi-C map and insulation profiles revealed that the insulation at element A is completely lost in ΔCTCF_A_, showing that CTCF acts as the main loop-extrusion barrier at element A, and the majority of A-anchored contacts (both putative E-P and E-E) were significantly weakened in this mutant, consistent with a role of CTCF and cohesin in their formation. Nevertheless, the decrease in A-anchored contacts was much smaller than in the ΔA mutant (compare **Fig. 5d, and Fig. 3d**) and the A-independent contacts remained unperturbed, strongly suggesting that the function of A as a scaffolding element cannot be explained by loop-extrusion (**Fig. 3a, 3d, 5a, and 5d**).

Together, these data strongly suggest that molecular attractions independent of loop-extrusion play an important role during differentiation-driven chromatin structure rewiring at the *Zfp608* locus.

### Looping interactions are crucial for rewiring during differentiation

In order to test the contribution of epigenetic interactions for rewiring, we generated 3D models of the *Zfp608* region using a polymer-based biophysical strategy based on Molecular Dynamics simulations (**Methods, and Extended Data Fig. 7a**). We represented chromatin as a chain of spherical beads, each containing 5 kb of DNA, featuring general biophysical properties including connectivity, excluding volume, and bending rigidity (**Methods**). This model was used to test three biophysical mechanisms (loop-extrusion, compartment interactions, and looping interactions) which can contribute to chromatin spatial organization (**Fig. 6a, and Methods**)^29–31^. We partitioned the *Zfp608* region into active (A) and inactive (B) compartments from previously published ESC and NPC Hi-C data and modelled attractions within and between them with energy strengths E_AA_, E_BB_, and E_AB_, which favour the formation of compartments through microphase separation^3,32–34^. Simultaneously, we added attractive looping interactions of strength E_L_ to model specific site-to-site contacts that were detected by the cHi-C (**Fig. 6a, 6b, Supplementary Table 5, and Methods**). In the same framework, we applied the loop-extrusion process that can form local domains and loops and where extruders, such as the cohesin complex, bind on chromatin at random positions and extrude loops in both directions until they unbind or encounter a barrier^27,28^. The extruder activity was characterized by the density (N_e_/Mb), lifetime (S_l_) on chromatin, and extrusion speed. Each barrier was described by its position on chromatin, direction, permeability (p), and pausing time (t_p_) induced on incoming extruders^35^. To limit the number of parameters to explore, we fixed the extrusion-speed to 1 kb/s and imposed that extruders can cross one another, consistent with recent works^36–38^. The barriers were positioned at the maxima of the experimentally determined insulation sites and they could stop extruders coming from both directions with the same permeability, but they were set to induce a direction-dependent pausing time (t_p_^L^ and t_p_^R^). To compare our models with the experimental data, we defined four metrics and tested more than 700 parameter sets, aiming to optimize these metrics (**Methods**)^39,40^. Similar to other studies, we found that more than one set of parameter values could similarly well describe the data. Hence, we visualized the metrics in a supervised selected set of best parameter values (**Supplementary Table 6**)^39,41^.

**Figure 6.**
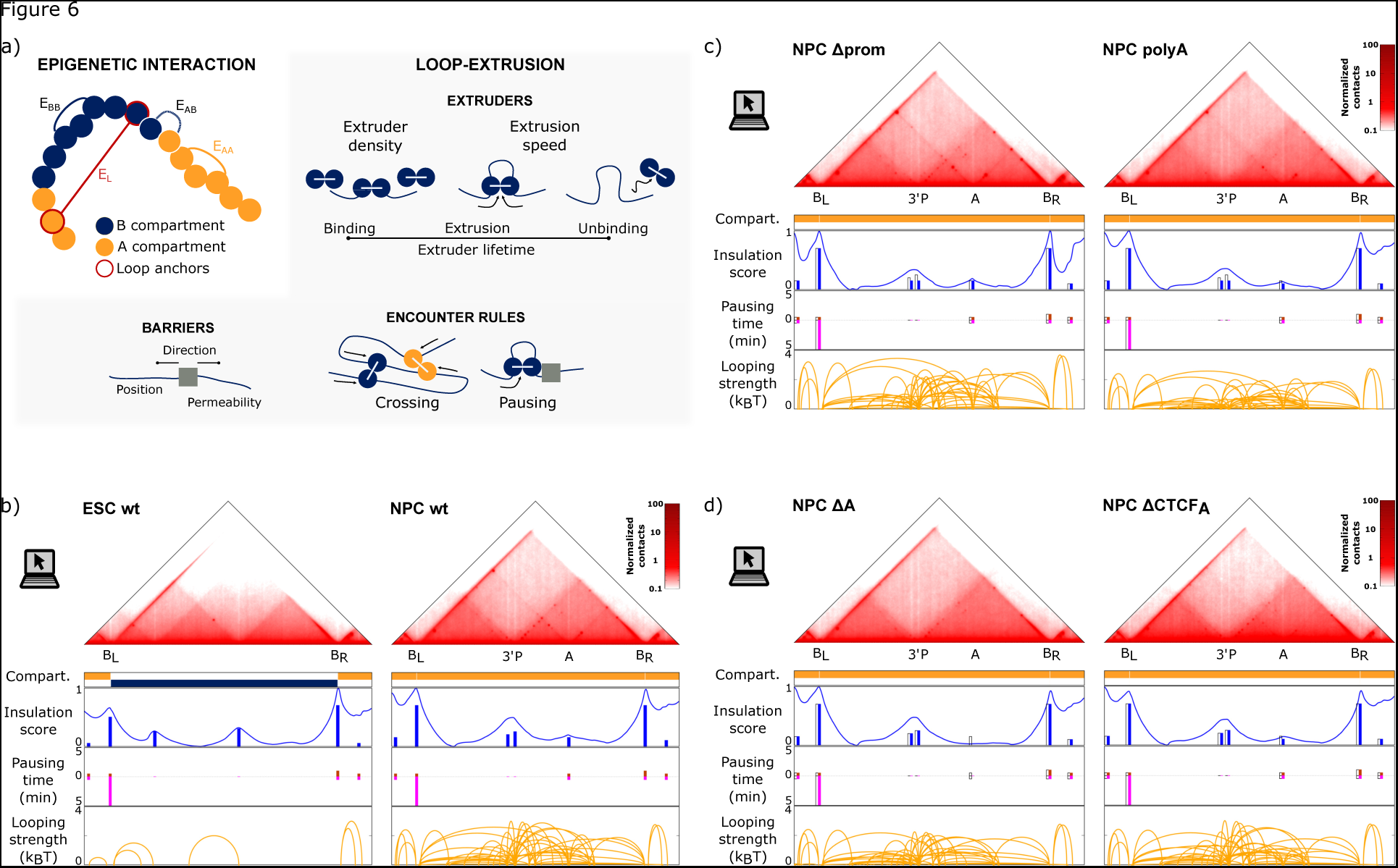
Biophysical models. **a)** Illustation of the biophysical mechanisms and parameters used for polymer modeling simulations. **b-d)** Normalized contact maps at 5kb resolution obtained from the optimal biophysical models generated for ESC and NPC wt (b), NPC Δprom and polyA (c), and NPC ΔA and ΔCTCF_A_ (d) are shown on the top (**Methods**). The four graphs below show the features of the experimental maps used as input for the simulations together with the parameters quantified using biophysical modeling. These include: the compartments partition inferred from the Hi-C dataset from Bonev et al.; the position of the loop-extrusion barriers marked as bars whose heights correspond to the optimized insulation strength1-permeability) together with the insulation score profile obtained from the models’ map; the left (positive values) and right (negative values) pausing times at each of the loop-extrusion barriers; and loops detected on cHi-C maps (**Methods**) shown as arcs whose height indicates the corresponding optimized attraction strength. In panels c) and d), barplots shown in colors are the permeablities and pausing times for the mutant, corresponding quantities in NPC wt condition are shown in white.

Using extensive data-driven parameter screening (**Methods, Extended Data** Fig. 7b-f**, and Supplementary Table 6**), we found that the optimized compartment-driven attractions were generally weak (0.00-0.02 k_B_T) and not compartment specific (E_AA_ = E_BB_ = E_AB_). In fact, models generated using compartment-specific attractions resulted in *cis*-decay curves (average number of contacts vs. genomic distance) that could not match the corresponding cHi-C decay at around 2.4 Mb (**Extended Data Fig. 7b, and c**). Furthermore, compartment-driven attractions higher than 0.02 k_B_T resulted in suboptimal models due to the excessive compaction of the region. We also ruled out extruder densities N_e_/Mb>8 and extruder lifetimes longer than 20 min, that induced an excessive compaction of the region between 100 kb and 1 Mb of genomic separation (**Extended Data Fig. 7d)**. In the optimized parameter sets, ESC and NPC wt models resulted in two similar loop-extrusion-dependent left and right barriers (B^L^ and B^R^) with high insulation strength (1-p=0.70-0.50) that generated a 2.4 Mb domain at the center of the region (**Fig. 6b**). The difference between the ESC and NPC state arose due to the internal barriers. In the NPCs, three novel, but weaker (1-p = 0.15-0.25) borders emerge between B^L^ and B^R^ at the enhancer A, the *Zfp608* promoter and the 3’ end of the *Zfp608* gene. Only combining these parameters with site-specific looping interactions (E_L_ = 3.00 k_B_T) resulted in models that optimally recapitulated experimentally determined contact patterns in ESC and NPC, indicating that loop-extrusion-independent molecular attractions are necessary to recover the chromatin organization observed at this locus *in vivo* (**Fig. 6b, Extended Data** Fig. 8a**, 8b, Supplementary Table 6, and Supplementary Video 1**).

We then identified the parameter changes that best explain the contact behaviour alterations in NPC mutant conditions compared to wt (**Methods**). The Δprom was best modelled by weakening the 3’ and *Zfp608* promoter barriers by ∼10% and by E_L_ increasing from 3 to 4 k_B_T in order to strengthen the *Zfp608* promoter-independent looping interactions **(Fig. 6c, and Extended Data Fig. 8c)**. These models confirmed that a partial loss of the promoter can impact not only insulation and contacts anchored at the deleted site but also strengthens promoter-independent contacts. The polyA condition was best captured by the same weakening of the barriers at 3’ and the *Zfp608* promoter in addition to a 5% weakening of the barrier at the A element and a 25% weakening of the promoter-anchored loops **(Fig. 6c, and Extended Data Fig. 8d**). These results suggest that transcription and its regulatory factors associated at the *Zfp608* promoter contribute to strengthening of the insulation at the *Zfp608* gene as well as to promoter-anchored looping interactions.

To account for the ΔA, ΔAB, and ΔB mutations, we removed the loop-extrusion (LE) barrier and the looping interactions involving the deleted elements (A and/or B). Notably, we found that these deletions also affected the insulation and the loops stemming from the *Zfp608* gene: in ΔAB and ΔB the insulation strength at the promoter decreased by 10 and 5%, respectively, and in ΔA and ΔAB the looping interactions from the promoter had to be reduced by 25% (**Fig. 6d, Extended Data Fig. 8e, 8f**, and **8g**). Simulating the ΔCTCF_A_ condition, we found that the LE barrier at element A still had a residual 10% insulation and the strength of the loops anchored in element A dropped by only 50%. We did not find any sizeable effect on the promoter for this mutant (**Fig. 6d, and Extended Data Fig. 8h**). The mild effect of ΔA, ΔB, and ΔAB on promoter-associated barrier permeability and looping interactions, that is not found in the ΔCTCF_A_ line, could therefore be due to binding of TFs other than CTCF at elements A and B.

These biophysical modelling results strongly suggested that loop-extrusion-independent molecular attractions could quantitatively explain chromatin conformation rewiring during differentiation. To better understand what might be driving these molecular attractions we investigated the factors bound at the anchors of NPC-specific loops. As expected, the anchors were acetylated in NPCs but also harboured PAX6 binding (**Fig. 1a, Extended Data Fig. 3a, and 3b**). As some of these elements are open and poised (harbouring H3K4me1 and H4K27Ac) already in ESC we focused on Pax6, a master regulator of NPCs that was proposed to play a role in NPC-specific contact formation (**Extended Data Fig. 3a, and 3b**)^3,40,42,43^. To this end, we used dCas9-VP64-p65-Hsf1 to ectopically overexpress *Pax6* in ESCs (ESC Pax6+) and performed RNA-seq, PAX6 ChIP-seq, and cHi-C in this mutant (**Extended Data Fig. 1i, 2e**, and **9).** We found that expression of multiple genes was changed between ESC Pax6+ and wt although the cells remained pluripotent (retained high *Oct4*, *Nanog*, *Klf4*, *Sox2, Esrrb,* and *Myc*) and lacked induction of expression of majority of the neuronal markers (i.e., *Nes, Ncam1, Msl1*, *Eno2*, *Foxa2*, *Dcx*) in agreement with previous studies **(Extended Data Fig. 2e, 9b, Supplementary Table 1, and 2)**^44^. However, *Pax6* overexpression did not induce transcriptional activity of *Zfp608* gene (**Extended Data Fig. 9a**). Moreover, the analysis of the PAX6 binding pattern in ESC Pax6+ cells showed that more than 36% of the PAX6-binding sites in ESC Pax6+ cells are much weaker or absent in NPCs, and vice versa (**Extended Data Fig. 9c).** In particular, PAX6 could not bind to any of its NPC-bound sites at the *Zfp608* locus except at the 3’ end of the gene (**Extended Data Fig. 9a**). Finally, we examined the cHi-C contact pattern which did not reveal major changes between ESC Pax6+ and ESC wt cells, indicating that chromatin conformation rewiring did not occur upon *Pax6* overexpression **(Extended Data Fig. 9a)**. These data show that Pax6 alone cannot act as a pioneer chromatin binding protein and cannot induce chromatin interactions at the *Zfp608* locus in ESCs, suggesting that critical cofactors required for its recruitment to its NPC-specific target sites might be missing.

## 4. Discussion

In this work we investigated chromatin rewiring during the ESC-to-NPC differentiation. Focusing on the *Zfp608* locus, we identified a distinct genetic element (element A) that directs this process. This is accomplished by scaffolding, mainly through a frequent/stable promoter-element A loop driven by multiple processes, including loop-extrusion and CTCF-independent chromatin looping that might depend on NPC-specific TFs and their cofactors.

We examined the contribution of multiple elements to rewiring by different mutants and showed that the deletion of element A induces the loss of most of the NPC-specific contacts, including A-independent contacts. This master role of element A suggests a hierarchical relationship between loops, in which A scaffolded loops promote the formation of additional contacts. Relatively little is known about the mutual relations between different kind of chromatin contacts in mammals. In *Drosophila,* one study found that so-called tethering elements form a subset of contacts early during early embryogenesis, before the establishment of topological domains. The emergence of these contacts fosters early, specific E-P loops that modulate the developmental genes’ activation kinetics^22^. A second study found extremely long-range contacts in fly neurons that can bridge distant domains and contribute to neuronal gene expression^7^. Similar extremely long-range chromatin contacts were also observed in mammalian neurons, suggesting that the hierarchical aspect of chromatin contacts might be present in other species^6^. In fact, earlier studies found that pre-formed chromatin topology strengthens the loop between the *Shh* promoter and the ZRS (*Shh* limb enhancer) thus providing transcriptional robustness during critical stages of development^20^. Our results extend these observations and suggest that there are multiple, distinct hierarchical layers of the E-P communication, and that it is likely that scaffolding elements, like A, are present at specific genomic loci to regulate chromatin architecture at different time points or in different tissues.

This scaffolding is directed by weak compartment-driven interactions, loop-extrusion and especially by site- and cell-state-specific looping interactions. Since the anchors of distinct loops are marked by H3K4me1 and H3K27Ac (promoter and element A) already in the ESC state, we speculate that these interactions are driven by NPC-specific TFs, perhaps including PAX6, a master regulator of NPC development, and its cofactors, rather than by H3K27Ac itself^42–45^. Other TFs have been previously demonstrated to participate in cell type specific looping and PAX6 has been found binding at the base of the NPC-specific loops^40,46,47^. Future experiments will be needed to assess directly if PAX6 contributes to looping at the *Zfp608* locus and if it requires additional factors for this function. Given that the human *Pax6* overexpression can induce ESC-to-NPC differentiation, but murine *Pax6* cannot, it is likely that there are inter-species differences for the PAX6 binding we might need to consider as well^44^.

Mechanistically, element A scaffolds the region through its frequent contact with the promoter which constrains the movement of adjacent chromatin regions and confines them to a closer space, thereby favouring the elements in the vicinity of A and the promoter to contact one another more frequently. Upon deletion of element A, chromatin becomes less constrained, in agreement with our cHi-C, 3D-FISH data and with recent simulations^48^. However, despite its enhancer function in cell culture assays, the deletion of element A did not cause significant aberrations to *Zfp608* expression. This finding suggests that one or several other enhancers which are present at the *Zfp608* locus might compensate for the loss of A, as observed previously in other systems^49–52^. However, it is remarkable to note that, even upon strong changes in the contact pattern in the whole region upstream to the *Zfp608* promoter, transcription is essentially stable. This suggests that *Zfp608* enhancers are able to correctly regulate transcription in the wake of substantial changes in 3D architecture, consistent with earlier work in which the effect of changes in enhancer-promoter contacts has been analysed and modelled quantitatively^53,54^. Interestingly, the deletion of tethering elements in flies similarly only causes a slight delay in transcriptional onset but this is sufficient to cause morphological changes later in development^22^. In mammals, a tight temporal regulation is imperative during *Hox* gene activation, where a deletion or insertion of CTCF sites delays or speeds up the activation of *Hox* genes, respectively^55,56^. Therefore, enhancer A might be either redundant with other elements or be specifically required for the regulation *Zfp608* expression in different cell types, timing, or in response to other cues^8,20,22^.

Other than structural elements driving local rewiring, we also demonstrated that transcription itself can modulate both insulation and local contacts. This finding is in agreement with other studies where genes involved in thymocyte differentiation were ectopically activated in ESCs, and their activation impacted insulation as well as the intra-gene contacts^57^. However, only a subset of ectopically activated genes (*Bcl6*, *Nfatc3*) changed local contacts and insulation and the ectopic activation of *Zfp608* in ESCs was unable to do so^3,57^. We suspect that this might be either due to the absence of NPC-specific TFs in ESCs or due to competition between the binding of dCas9 with TFs needed for chromatin rewiring in ESCs. However, we demonstrated that in the polyA mutant in NPC, in which there is no such binding obstruction at the promoter, transcription weakening is sufficient to diminish insulation and contacts. These data provide evidence for a role of transcription to establish local chromatin conformation (insulation and contacts) upon native transcriptional induction. What remains to be understood are the relative contribution of RNA polymerase and its associated cofactors versus that of TF binding and histone marks at the promoter. Furthermore, CTCF/RAD21 binding at the *Zfp608* promoter might contribute to regulate the frequency and/or the duration of promoter-anchored contacts in conjunction with the transcription^17,18^. Future work will be required in order to test this possibility and to disentangle the respective role of different components in the multi-layered regulation of chromatin architecture.

## 5. Material and Methods

### EXTENDED METHODS

#### EXPERIMENTAL MODEL AND SUBJECT DETAILS

##### Cell Lines

E14GT2a p14 cells were purchased from MMRRC, UC Davis and used as ESC wt condition and for the generation of all here-engineered cell lines described below. ESC cells were cultured in a high-glucose GMEM media (ThermoFisher #21969035) supplemented with 1x GlutamaX (ThermoFisher #35050038), 1x Penicillin/Streptomycin (ThermoFisher #10378016), 1x sodium pyruvate (ThermoFisher #11360070), 1x non-essential amino acids (ThermoFisher #11140035), 0.1 mM 2-mercaptoethanol (ThermoFisher #31350010), ESC-grade tetracycline-free FBS (ThermoFisher #26140079) and 1000 U/mL LIF (Millipore #ESG1107). NPC differentiation was done using 4.1 million ESCs per experiment using a previously published retinoic acid-based protocol^58^. During differentiation, cells were cultured for the first four days in high glucose DMEM (ThermoFisher #21710025) supplemented with 1x Glutamax (ThermoFisher #35050038), 1x non-essential amino acids (ThermoFisher #11140035), 0.1 mM 2-mercaptoethanol (ThermoFisher #31350010), 1x Penicillin/Streptomycin (ThermoFisher #10378016) and tetracycline-free FBS (GE Life Sciences HyClone #SH30071.03). The media was changed every two days and on day four the media was additionally supplemented with 5 µM retinoic acid. Cells were kept in retinoic-acid-supplemented media until day 8 with media changes every two days.

##### Generation of CRISPR-Cas9 and CRISPR-dCas9 Lines

gRNAs were designed to achieve the deletion of element A, element B, a part of the *Zfp608* promoter, to destroy a CTCF binding site at element A, and to make a single cut for the polyA insertion. The gRNAs were predesigned with a BbsI or a SapI overhang to be ready for the subsequent insertion into the px459 plasmid (addgene #62988). The pairs of gRNA sequences for deletions were as follows: for ΔA (gRNA1: [PHO]**CACC**GGGCCACCATTAACCTTTGT and gRNA2: [PHO]**CACC**GTGCACGGGAAACCCTTCAGA), for ΔB (gRNA1: [PHO]**CACC**GCACACTTGCAATACGGTCAT and gRNA2: [PHO]**CACC**GTGCTTGATTCTAGTCGAAGA), for Δprom (gRNA1: [PHO]**CACC**GCAAGCCTTTCACATCACGTG and gRNA2: [PHO]**CACC**GATGTGAGGATCCGACCAAGG); and two single gRNAs for CTCF motif destruction (gRNA: [PHO]**CACC**GTCTGCTGGCTGATGTTCCAA) and for the polyA insertion (gRNA: [PHO]**CTC**GTTCACAACCCGAGCACCGA). For the deletion constructs gRNAs were cloned into px459 plasmid (addgene #62988) carrying a different resistance marker so that one gRNA is in a plasmid with Puromycin and the other one in a plasmid with Neomycin resistance gene. For the polyA insertion, an N1 donor plasmid was constructed carrying two homology arms adjacent to gRNA cut site (centromeric arm (750 bp) and telomeric arm (1500 bp)) that flanked 3x SV40 polyA signal. Furthermore, in the donor plasmid’s homology arm the PAM sequence was mutated so that the gRNA would not re-target it once the construct was inserted.

The E14 wt cells were transfected with Lipofectamine 3000 (ThermoFisher #L3000001) in standard ESC media (composition above) without Penicillin/Streptomycin and left over night. Subsequently, the cells were selected either with Puromycin only (2 µg/mL) (if carrying one gRNA) or with Puromycin (2 µg/mL) and Neomycin (1000 µg/mL) if carrying two plasmids (two gRNA, or one gRNA and an N1 donor plasmid) for four days. Afterwards, cells were allowed to recover for three to four days and clones were picked and genotyped (directly in a 96-well-plate replica of picked clones – using ThermoFisher Phire kit #F170S according to the manufacturer’s recommendation). Selected clones were expanded, sequenced and then the breakpoints were amplified and subcloned into a TA cloning vector and sequenced to confirm that the breakpoints were present on both alleles.

*Pax6* gRNA (CTCGGCCTCATTTCCCGCTC) was cloned previously in the laboratory into the Bleomycin resistant lentiviral backbone plasmid (addgene #61427) and made into a virus. Then, the ESC dCas9-VP64/p65-Hsf1 cells (from Bonev et al.) were transduced with the *Pax6* gRNA carrying lentivirus (with BleoR) using a standard ESC media without Penicillin/Streptomycin and supplemented with polybrene (8 μg/mL) for one day^3^. Subsequently, the cells were treated with Zeomycin (500 μg/mL) for two weeks and left to recover.

#### EXTENDED METHOD DETAILS

##### Design of cHi-C probes

Capture Hi-C probes were designed using the a previously published cHi-C design tool GOPHER and Agilent SureSelect wizard (https://earray.chem.agilent.com/suredesign/home.htm)^59^. The entire region for the capture (mm10, chr18:52,439,203-57,439,739) was loaded into the GOPHER tool and the capture region was divided into the series of regions that were maximum 250 pb away from the nearest DpnII site, not at the DpnII clusters or the repetitive regions and have more than 80% GC coverage. It was made sure these regions tile the entire capture region but that they do not overlap as not to skew the coverage of the region. Then, the pre-select regions from the GOPHER output were loaded into the Agilent SureSelect wizard which designed a 120 bp long RNA probes using moderately stringent masking, balanced boosting and 3x tiling. The GOPHER output that can be used for Agilent SureSelect input is provided in **Supplementary Table 3** and the final SureSelect Agilent probes are reachable with Agilent France under the SureSelect capture kit #5190-4816 with specific probe design #3132861.

##### Cell Isolation, Purification and cHi-C

ESC and NPC cells were collected with Tryple (ThermoFisher #12604013), fixed with 1% methanol-free formaldehyde in ESC or NPC media and incubated for 10 minutes with rotation at room temperature (RT). After, cell lysis, digestion, biotin fill-in, ligation, and sonication were performed as previously published (Rao et al. 2014)^60^. End-repair, biotin pulldown, polyA tailing, adaptor ligation, pre-capture PCR, capture, library preparation and post-capture PCR were performed according to manufacturer’s recommendations using the Agilent capture kit (Agilent #5190-4816 with the SureSelect design #3132861), library preparation (Agilent #G9611A) and Herculase polymerase (Agilent #600677) with slight modifications listed hereafter (https://www.agilent.com/cs/library/usermanuals/public/G7530-90000.pdf). Modifications: After the biotin pulldown, the polyA mix (prepared as per Agilent protocol) was added directly onto the streptavidin beads with from the biotin pulldown and incubated for 30 minutes at 37 °C. At the end of the reaction the beads were pulled with the magnet, the supernatant was discarded and beads were washed with 100 μL of water. Then, the adaptor ligation mix (prepared as per Agilent protocol) was added directly onto the streptavidin beads and incubated for 15 minutes at 20°C. After the reaction was done, the beads were separated on the magnet, the supernatant discarded and beads washed with 100 μL water. Finally, the pre-capture PCR mix (prepared as per Agilent protocol) was added directly onto the beads and each sample was split into four 25 μL PCR reactions. After the PCR, the supernatant was purified and used for capture reaction and the beads were washed and stored in 50 μL water in -20°C as backup. The final library (post-capture PCR) was purified with SPRI beads (Beckman Coulter #B23318) with two-sided selection in the ratios 0.5x and 0.8x. Final libraries were sent to BGI China where they were sequenced with 150 bp paired-end reads.

##### RNA isolation and RNA-seq

Cells were collected with Tryple (ThermoFisher #12604013), lysed and RNA was isolated using RNAeasy mini kit (Qiagen #74104) per manufacturer’s recommendations. Additional, on-column DNAseI digestion (Qiagen #79254) was performed as recommended. The RNA purity and integrity was checked on a gel and the RNA was sent to Novogene where rRNA depletion was performed and libraries were constructed. Samples were sequenced using 150 bp paired-end reads.

##### Quantitative real-time PCR (qPCR)

qPCR was performed on cDNA generated with 1 µg of RNA with Maxima First Strand cDNA Synthesis Kit (ThermoFisher #K1672) according to the manufacturer’s recommendation. Then, the above generated 20 µL of cDNA was diluted with 80 µL of water and immediately used as template. The qPCR primers were constructed in several exons, exon-exon spanning, introns or exon-intron spanning in *Zfp608* gene as follows: (i) EXONS: e1-e2 (F-AAGTAAAATGAGGCACAGGGCT and R-TCGAGGTCCAGCTTCTCCTTT), e3 (F-CCCAAGACACTCTGGCTCTG and R-AGCACCAGTCGAACAGCTTT), e4-e5 (F-CCTCCTAGACTGCACCAAGC and R-GACCTCGCTCTCTTCCCTCT), e5 (F-GATGGGGAACTGGCCTTTGA and R-CAGCAGCTAAGACCAGCAGT), and e6 (F-CTGGACACGCCACTCTCTTT and R-ACCTGGACCAACAAAAGCCA) ; (ii) INTRONS and INTRON SPANING: i2a (F-CTAGCTTCCTAAACAGGCATCC and R-TGGCATGATAGCACCCAGAG), i2b (F-GGGTAGCCCAATGTTTCCTTAG and R-TTGCCAGCGATCTGAACATC), e4-i4 (R-AACAGGATGGGAGGAGCAGA and R-ACCCTCCTAGACTGCACCAA) and i4 (F-CTGGTGGCTTGAGAGTGAGG and R-ACATAATCCACAGGGCAGCC). The qPCR was done using LightCycler 480 Sybr green master mix (Roche #04707516001) with 10 µl individual reactions in technical triplicates and biological duplicates according to the manufacturer’s recommendations.

##### ChIP-seq

Cells were collected with Tryple (ThermoFisher # 12604013) and fixed with 1% methanol-free formaldehyde in ESC or NPC media and incubated for 10 minutes with rotation at RT. For every transcription factor ChIP, 4 million cells were used and for every histone modification ChIP, 2 million cells were used. ChIP-seq was done using Diagenode iDeal ChIP-seq kit for transcription factors (#C01010172) or histones (#C01010171) following the manufacturer’s protocols. The antibodies used for the ChIP-seq were as follows: H3K4me3 (Millipore #04-745), H3K27meAc (Active Motif #39133), H3K36me3 (abcam ab9050), PolII (Diagenode #C15100055), CTCF (Active Motif #61311), RAD21 (abcam ab217678) and PAX6 (Millipore AB2237).

##### Oligopaint probe design and preparation

Oligopaint libraries were constructed following the procedures described by Beliveau *et al.*, see the Oligopaints website (https://oligopaints.hms.harvard.edu/) for further details. Libraries were synthesized by GenScript in the 12K Oligo pool format^61^. The coordinates (mm10), size, number, density of probes and primers used for the libraries are listed in **Supplementary Table 4.**

Oligopaint libraries were designed using the mm10 Harvard ‘balance’ BED files, which consist of 35–41-bp genomic sequences throughout the regions of interest. BED files can be retrieved from the Oligopaint Harvard website. Each library contains a universal primer pair followed by a specific primer pair (barcode without the ‘Sec’ sequences) hooked to the genomic sequences (117–125 bp in total). Oligopaint libraries cover 10 to 15 kb of the genomic sequence of interest (i.e., the *Zfp608* promoter, the A and the B elements), representing 100–150 oligonucleotides distributed along each specific region. Oligopaint libraries were produced by emulsion PCR amplification from the oligonucleotide pool followed by a ‘two-step PCR’ procedure and digestion by lambda exonuclease^61^. Here, to increase the sensitivity of the probes, we developed a ‘double-tail’ strategy (dt). The first PCR (PCR1) produces a PCR fragment containing the genomic sequence flanked by two secondary oligonucleotide binding sites (Sec2/Sec1 or Sec4/Sec6 or Sec3/Sec5 binding site pairs) for signal amplification. The second PCR (PCR2) hooks a Phosphate atom at the 5’-end and a fluorophore (Alexa Fluor 488, ATTO 565 or ATTO 647) at the 5’-end on the opposite strand. The lambda exonuclease digests the PCR product from the 5’P, leading to the production of pool of single strand primary oligos bearing one fluorophore on their 5’-end. This probe can be used for FISH together with secondary oligos that are dual labelled at their 5’- and 3’-end, complementary to their binding sites (Sec2/Sec1 or Sec4/Sec6 or Sec3/Sec5) and bringing additional 4 fluorophores per oligo, each oligo carrying five fluorophores in total (Alexa Fluor 488, ATTO 565 or ATTO 647 fluorophores). This new strategy allows us to produce reliable 10 kb Oligopaint probes with good confocal-based sensitivity. All oligonucleotides used for Oligopaint production were purchased from Integrated DNA Technologies. Oligonucleotide primer sequences (5ʹ→3ʹ) used in this study are listed in Supplementary Table 4.

##### FISH procedure

The FISH protocol is described in Szabo et al.^62^. Briefly, ESCs were grown directly on coverslips (170 ± 5 µm (ZEISS) in 6-well plates for 2 days. For NPCs, at the end of differentiation, aggregates were dissociated with Tryple (ThermoFisher #12604013) and resuspended at a concentration of 1.5 × 10^6^ cells/ml in DDM media (DMEM/F12 (ThermoFisher #31331-028), 500 µg/mL BSA (ThermoFisher #15260037), 1x non-essential amino acids (ThermofFisher #11140035), 1x Penicillin/Streptomycin (ThermoFisher #10378016), 1x sodium pyruvate (ThermoFisher #11360070), 0.1 mM 2-mercaptoethanol (ThermoFisher #31350010) and 1x N2 supplement (ThermoFisher #17502048)). Then, the single cell suspension was plated onto the coverslips pre-coated with 33 µg/ml of poly-l-lysine in DPBS (SigmaAldrich #P8920) and 3 µg/mL of laminin in DPBS (SigmaAldrich #11243217001) in 6-well plates for 2 hours at 37 °C. Cells were washed once in PBS and fixed for 10 min in 4% PFA in PBS, rinsed with PBS, permeabilized for 10 min in 0.5% Triton X-100 in PBS, rinsed with PBS, incubated for 10 min in 0.1 M of HCl and rinsed with 2× SSC/0.1% Tween 20 (2× SSCT). Cells were then incubated for 20 min in 50% formamide/2× SSCT at RT followed by 20 min in 50% formamide/2× SSCT at 60 °C. The 1–3 µM final concentration Oligopaint probe mixtures contained the same amount of their secondary oligos and 0.8 µl of RNase A (10 mg/mL) in 20 µl of FISH hybridization buffer (FHB: 50% formamide, 10% dextran sulfate, 2× SSC and salmon sperm DNA (0.5 mg/ml)). For good homogenization in FHB, probe mixtures were incubated on a thermomixer for 10 min at 80°C at 1500 rmp and let cool down to RT before use. Probe mixtures were added directly to coverslips that were then sealed on glass slides with rubber cement (Fixogum; Marabu). Cellular DNA was co-denatured with the probe mixture for 3 min at 80 °C on a heating block immersed in a water bath and hybridization was performed overnight (16 to 20 hours) at 42 °C in a dark and humid chamber. Cells were then washed for 15 min in 2× SSCT at 60 °C, 10 min in 2× SSCT at RT, 10 min in 0.2× SSC and twice in PBS. Cells were then incubated for 10 min in PBS with DAPI at RT (final concentration at 0.2 to 0.5 µg/ml) and washed twice for 5 min with PBS. Coverslips were mounted on slides with VECTASHIELD (CliniSciences) and sealed with nail polish.

##### Image acquisition

Confocal imaging was performed using a Confocal Zeiss LSM980 Airyscan II equipped with a x63/1.4 numerical aperture (NA) Plan Apochromat oil immersion objective. Diodes laser 405, 488, 561 and 639 nm were used for fluorophore excitations, leading to blue, green, red and far-red channels. Acquisitions were performed in 3D using the AiryScan Multiplex 4Y mode, with a pixel size of 40 nm and a z-axis step of 140 nm. 15-20 positions were scanned per FISH slide, and z-stacks were mounted in independent channels for analysis using the FIJI software^63^. Each experimental condition was analyzed in triplicate (from three independent NPC differentiation). Replicates were merged for data analysis leading to 40-50 z-stacks per condition.

##### Image analysis

The 3D distance measurement in 3 colors was adapted from Szabo et al.^62^. 3D FISH analysis was conducted using the “Image Processing Toolbox” in MATLAB vR2023a. Briefly, channels were smoothed using a Gaussian filter (σ = 3 and 1 pixel for DAPI and FISH channels, respectively) and segmented in 3D using an adaptative threshold; FISH objects that were smaller than 900 voxels or located outside of the DAPI-segmented regions were discarded. 3D distances between the centroids of segmented FISH objects were calculated, and for the three objects (from the green, red and far-red channels), only the mutual nearest neighbors that were closer than 1.5 µm were considered. In each individual nucleus, the three distances were paired allowing for triangle area measurements and optimal superimposition and kernel density analysis.

##### Superimposition of FISH data by Optimal Superimposition

To examine the relative positions of the three loci (promoter, element A, and element B) in space, we devised a novel method inspired by geometric morphometrics. Since the three loci form a triangle in space, we performed an optimal superimposition to align the triangles, without scaling, over their barycenters using the R package ’paleomorph’ with default parameters. The triangle’s shape causes all its vertices to align within a 2D plane (**Extended Data Fig. 6d**). To enhance visualization, an orthogonal projection was performed using PCA of the optimal superimposition (**Extended Data Fig. 6d**)^64^. This process ensured that the original data’s positional identity and distances were maintained despite the alignment. Kernel density calculations were employed to examine the spatial occupancy of each locus. Comparison between conditions was conducted by evaluating the area at a specific kernel density across various density levels.

#### QUANTIFICATION AND STATISTICAL ANALYSES

##### ChIP-Seq analysis

ChIP-seq samples were mapped using bowtie2 v.2.3.5.1 (https://bowtie-bio.sourceforge.net/bowtie2/index.shtml) with command “*bowtie2 -p 12 --no-mixed --no-discordan*t” ^65^. Then, we used samtools v.1.9 (https://www.htslib.org/doc/samtools-view.html) to filter out low-quality reads (command “*samtools view -b -q 30*“), to sort the bam files (command *“samtools sort”*), and index them (*“samtools index”*) with default parameters^66,67^. Afterwards, bigwig files were produced using the deepTools package (https://deeptools.readthedocs.io/en/develop/content/tools/bamCoverage.html) with command “*bamCoverage --normalizeUsing RPKM --ignoreDuplicates -e 0 -bs 10*” to make sure that the resulting bigwig files are normalized and without duplictes^68^ . Finally, the ChIP-seq tracks in **Fig. 1b, 2a, 2b, 3b, 5a, 5b, Extended Data** Fig. 1b-i**, 5b, and 9a** were visualized using the IGV v.2.16.1 software^69^. For the PAX6 ChIP-seq we called peaks with MACS2 (https://hbctraining.github.io/Intro-to-ChIPseq-flipped/lessons/06_peak_calling_macs.html) with a q value cutoff 0.01 in ESC wt and NPC wt^70^. The summits of both peak sets were extended 150 bp left and merged if closer than 300 bp. Then, *k-means* analysis was applied using *in-house* R scripting where to cluster the merged peak set using the average signal at the peak summit +/- 250 bp in the bigwig files of each PAX6 ChIP-seq (in ESC or NPC) or with ATAC-seq (GSE161993) (**Extended Data** Fig. 9c**, and d**).

##### RNA-seq analysis

RNA-seq samples were mapped using STAR aligner v2.7 (https://github.com/alexdobin/STAR/blob/master/doc/STARmanual.pdf) and the bigwigs were produced and visualized as for the ChIP-seq samples (**Fig. 1b, 2a, 2b, 3b, 5a, 5b, Extended Data** Fig. 1b-i**, 4a, 4d, 5b, and 9a**)^71^. Additionally, Subread v2.0.6. (https://subread.sourceforge.net/) was used (command “*featureCounts -p –countReadPairs -s 2*”) to generate the count table (**Supplementary Table 2**)^72^, that was used as an input for the DEseq2 (https://bioconductor.org/packages/devel/bioc/vignettes/DESeq2/inst/doc/DESeq2.html) to compute the FPKM values (**Supplementary Table 1**) and to perform the differential analysis (**Supplementary Table 2**)^73^. Additionally, we generated FPKM tables and performed differential analysis for two feature files: 1) UCSC RefSeq GTF file (https://hgdownload2.soe.ucsc.edu/goldenPath/mm10/bigZips/genes/mm10.refGene.gtf.gz) and 2) a custom GTF file (the UCSC RefSeq GTF + manually added *Zfp608A* gene where we only the introns of the *Zfp608* gene (**Supplementary Table 1, and 2**). This way we were able to quantify FPKM and normalized counts in exons and introns of *Zfp608* gene shown in **Fig. 2c, 3c, Extended Data** Fig. 4c, 4f, 5e, and **9b.**

##### cHi-C data analysis

###### Generation of the cHi-C matrices

Capture HiC samples were analyzed using the TADbit pipeline^74^ (https://github.com/3DGenomes/tadbit) which was used to (i) check the quality of the FASTQ files; (ii) map the paired-end reads to the *M. musculus* reference genome (release *mm10* downloaded from http://igenomes.illumina.com.s3-website-us-east-1.amazonaws.com/Mus_musculus/UCSC/mm10/Mus_musculus_UCSC_mm10.tar.gz) using bowtie2 v.2.3.5.1 (https://bowtie-bio.sourceforge.net/bowtie2/index.shtml) with command line “*bowtie2 -x bowtie2Index -p 8 --reorder -k 1*” accounting for the restriction-enzyme (*DpnII*) re-ligation-sites (*fragment-based mapping)* ^65^; (iii) remove non-informative reads using the default TADbit filtering options ^74,75^. Differences between replicates (2 per each of the 11 conditions) were measured using the <u>S</u>tratum- adjusted <u>C</u>orrelation <u>C</u>oefficient (SCC) applying *HiCRep* v1.12. (https://github.com/TaoYang-dev/hicrep) on the capture region (chr18:52,439,203-57,439,739) using the *get.scc* function with parameters resol=20kb, lbr=0, ubr=2000000, h=1 which was previously trained using the *htrain()* on the NPC wild type replicates^76^. Using 1.0-SCC, as a measure of the similarity (0 - similar and 1 dissimilar) between replicates and hierarchical clustering analysis using *hclust()* function in R with Ward.D2 method allowed to distinguish and group together the replicates of the different conditions (**Extended Data** Fig. 1a) motivating us to merge the valid-pairs of different replicates in a unique dataset per each condition. The total numbers of sequenced reads, uniquely mapped reads, and valid read-pairs per sample are reported in **Supplementary Table 7**. The numbers of contacts within the capture region (cisContacts), between the capture region and chromosome 18 (transContacts), and between regions outside the capture region (outContacts) were measured with in-house bash scripts, and reported in **Supplementary Table 7**. .cool files were generated using cooler v.0.9.1 (https://cooler.readthedocs.io/en/latest/index.html) from the *cisContacts* only at 100 bp resolution (command “*cooler cload pairs --assembly mm10 --chrom1 1 --pos1 2 --chrom2 6 --pos2 12 ./scripts/chrom_sizes.txt:100*”)^77^. *.mcool* files at various resolutions 2, 5, 10, 15 and 20 kb were obtained and normalized via the Iterative <u>C</u>orrection and <u>E</u>igenvector decomposition algorithm (ICE) with default parameters (command “*cooler zoomify -r 2000,5000,10000,15000,20000,40000 file.cool -o file.mcool --balance*”)^78^. Applying the iterative correction normalization algorithm only to the capture region ensured a seamless convergence of the method for all samples. When comparing architectural features between different conditions, down- sampled datasets have been obtained by randomly selecting a number of *cisContacts* equal to the minimum within the compared datasets. Namely, ESC samples in **Extended Data** Fig. 1c-i**, and 9a** were down-sampled to ESC wt dataset (4,043,665 *cis-contacts*) and NPC samples in **Fig. 2a, 3a, 5a, and Extended Data** Fig. 5a to the NPC wt dataset (6,596,808 *cis-contacts*). Map resolution of 5 kb for the entire cHi-C dataset (including down-sampled and single-replicate samples) has been defined as the smallest locus size such that 90% of loci in the entire capture region have at least 1,000 contacts. The map resolution is meant to reflect the finest scale at which one can discern architectural features reliably^60^. Valid interactions were stored in a database using the “misha” R package (https://github.com/msauria/misha-package). The “shaman” R package [https://bitbucket.org/tanaylab/shaman] has been used for computing the cHi-C expected models with parameters grid_small=0.5e6 bp and grid_high=1e6 bp using the function shaman_shuffle_hic_track() and cHi-C scores with parameters k=250 and k_exp=500 using the function shaman_score_hic_mat() (**Fig. 1a, 2a, 3a, 5a, Extended Data** Fig. 1b-i**, 5a, and 9a**). The maps are represented with a colour scale from - 100 (blue – depleted contacts with reference to the expected model) to 100 (red – enriched contacts with reference to the expected model) showing the KNN normalized score (K-number (250) nearest neighbor normalized).

**Insulation score analysis**. The insulation was computed on the observed cHi-C dataset binned at 2 and 5 kb resolution with windows of 300, 350, 400, 450, and 500kb using the function insulation() of the *cooler* package (**Fig. 1b, 2a, 3b, 5a**, and **Extended Data** Fig. 5b)^77,79^. The quantification of the insulation scores (IS) at regions of interest was performed on the IS binned at 2 kb resolution for the five considered window values (300, 350, 400, 450, and 500 kb) contained in each region of interest. The IS normalized between 0 (absence of insulation) and 1 (maximum insulation). The pairwise statistical comparison of the IS quantifications, including the p*-values* reported in **Fig. 2a, 3b, and 5a** resulted from a Welch t-test (H0: true difference in means is equal to 0. The variances of the samples are thought not to be equal) between the NPC wt condition and each of the mutants.

###### Analysis of looping interactions

Loops were called using mustache v1.0. with default parameters ICE-balanced maps at 5, 10, and 15 kb resolution for all conditions at 5, 10, and 20 kb on ICE-balanced maps^80^. Next, the identified set of loops called at each resolution were merged and filtered applying the following criteria:

- All loops with *fdr* >= 0.20 were removed.
- All loops whose anchors were closer than 40 kb were removed.
- All loops where at least one anchor was closer than 40 kb from a marginal bin as defined in *cooler*. At a given resolution, a marginal is a bin of the interaction matrix which is removed from the ICE-normalization using the MAD-MAX procedure because it has so few contacts that may affect the convergence of the normalization procedure^77^. We notice that in each of the cHi-C dataset, all the genomic regions outside the capture region are marginals. Hence, this filter excludes any loop that is too close to the limits of the capture region.
- Two loops whose both anchors are closer than the resolution are merged in a single loop.

The sets of filtered and merged loops are further subjected to manual scrutiny in *higlass* (http://higlass.io/) to remove any spurious detection and/or missing looping interaction^81^. The final sets of loops in ESC wt and NPC wt are reported in **Supplementary Table 8.** Next, for each detected loop, we considered 10 kb-regions around the central nucleotide of each of the anchors and measured the corresponding number of observed interactions in each of the cHi-C replicates. This matrix of interaction counts was used as input of the Deseq2 software v1.40.2 (https://bioconductor.org/packages/devel/bioc/vignettes/DESeq2/inst/doc/DESeq2.html)to quantify differential loops interactions between the NPC wt condition and ESC wt, or NPC mutants (**Fig. 1d, 2e, 2f, 3c, 3d, 5d, and Extended Data** Fig. 5d,)^73^. p-adjusted values were calculated using Wald test automatically with DEseq2.

##### Chromosome polymer model

Two polymeric systems were prepared in 50 replicates each to model 1 copy of the chromosome region chr18:53,500,000-56,900,000 bp (*Zfp608* region) in ESC and NPC state, respectively. Each *Zfp608* region was represented as a chain of beads where each monomer of unitary mass (m = 1.0) hosts 5 kilobase pairs (kbp) of DNA sequence and has a diameter of σ. This representation was obtained with the Kremer-Grest bead-spring model with the same parameter as Di Stefano et al.^82,83^.

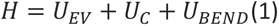

The first term was a truncated and shifted Lennard-Jones potential that controls the *cis*- and *trans*- chromosome excluded volume interactions:

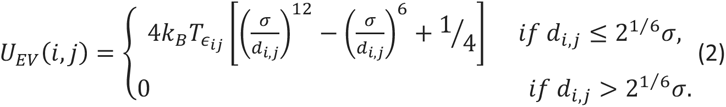

where k_B_ is the Boltzmann constant, T the temperature, *ε*_ij_ is equal to 10 if |*i* – *j*| = 1, and 1 otherwise, σ was the thickness of the chain and d_i,j_ is the modulus of 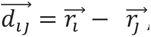, that is the distance vector between the monomers *i* and *j* at positions *r_i_* and *r_j_*, respectively.

The second term was a FENE potential that ensures the chain connectivity between consecutive beads on the same polymer chain:

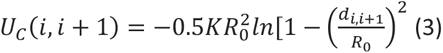

where K=0.33 k_B_T/nm^2^ and R_0_=1.5σ. The combined action of the connectivity and excluded volume interaction between consecutive beads was such that the average bond length was close to σ and never exceeded 1.1σ.

The third term is a (Kratky-Porod) bending potential:

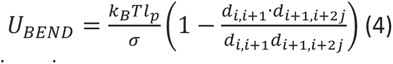

where l_p_ is the chain persistence length.

The dynamics of the polymer model was simulated using the LAMMPS simulation package (version 29 Oct 2020) integrating the (underdamped) Langevin equation of motion^84^:

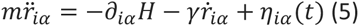

where m is the mass of the bead that was set equal to the LAMMPS default value, H is the energy of the system in **Eq. (1)**, the index *i* runs over all the particles in the system, and *α* = (*x*,*y*,*z*) indicates the Cartesian components, and γ = 0.5 τ^−1^ is the friction coefficient with τ = σ(m/ε)^1/2^ is the Lennard-Jones time. The stochastic term η_iα_ satisfies the fluctuation-dissipation conditions. The integration time step used in the numerical integration was equal to Δt = α τ_LJ_, where the factor α was adapted, as specified below, to the different stages of the preparation and production runs.

###### Preparation of the initial conformations

Each chain is initially organized in a rod-like folding featuring rosettes along the main axis, and placed in random positions inside a cubic simulation box of side 600σ with periodic boundary conditions, avoiding clashes with other chains^39,85^. After an energy minimization (*LAMMPS command*: minimize 1.0e-4 1.0e-6 100000 100000), each of the polymeric system is compressed to reach the target DNA (ρ) and volumic densities (ɸ) of ESC, ρ_ESC_∼0.006bp/nm^3^ and ɸ_ESC_∼3%, and NPC, ρ_NPC_∼0.011bp/nm^3^ and ɸ_NPC_∼5%, nuclei respectively. These estimates were done by considering the chromosomal DNA content of mouse cells (chr1-19, X, Y, M for diploid cells) of DNA=5.45 × 10^9^ bp, nuclear volumes of V_ESC_∼1000 μm^3^ and V_NPC_∼522 μm^3^, nucleolar volumes v_ESC_=130 μm^3^ and v_NPC_∼3 μm^3^ ^86–88^, chromatin diameter D_c_=0.014 μm by assuming a chromatin clutches model with σ = lp = 14 nm , every 1kb of chromatin^89,90^. Assuming that chromatin binding proteins increase chromosome fiber volume by 2. Finally, we assumed that the mutations (NPC ΔA, NPC ΔAB, NPC ΔB, NPC Δprom, NPC polyA, NPC ΔCTCF) don’t introduce sizable variations of these wt parameters. These conditions were achieved by molecular dynamics simulations (LAMMPS command fix 1 all nph iso *P P* 2.0, where P_ESC_=0.23 and P_NPC_=0.44) of 10 τ_LJ_ (10,000 Δt with Δt = 0.001 τ_LJ_). At the target densities, the polymer chains have parameters σ_ESC_∼36nm and l_p,ESC_∼54nm for ESC and σ_NPC_∼35nm and l_p,NPC_∼56nm for NPC^91^. Finally, each polymeric system is relaxed with molecular dynamics run of 60,000 τ_LJ_ (5,000,000 Δt with Δt = 0.012 τ_LJ_). By comparing the average monomer <u>M</u>ean-<u>S</u>quared <u>D</u>isplacement (MSD) in these relaxation runs and the MSD of non-transcribed genes measured in Gu et al. by live-cell imaging, we provide an estimate of the correspondence between the simulated time (in τ_LJ_) and the timescale of seconds (**Extended Data** Fig. 7)^92^. This analysis revealed that ∼15 τ_LJ_ and ∼18.6 τ_LJ_ of simulation map onto 1s in the ESC and NPC system respectively (**Extended Data** Fig. 7a). The conformations after 1,200 τ_LJ_ are next used as the initial conformations for the downstream simulations.

###### Biophysical modeling

Using the data generated in this work, we informed and parameterized three distinct mechanisms: compartments interactions, loop-extrusion, and site-specific looping shape the structural organization of the *Zfp608* region^27,31,35,93,94^. In the following, we detail the parameters involved in each mechanism, the strategy to parameterize the models, and the downstream data analysis of the obtained structures.

*Model parameters*. Short-range interactions were used to test the attractions between the model regions which correspond to A/B compartments as identified in Bonev et al. and the site-specific chromatin looping^3^. These interactions have been modeled using attractive Lennard-Jones potentials (see **Equation 2**) with cutoff=2.5σ, that allowed to include the attractive part of the Lennard-Jones potential. The strengths of compartments’ interactions (ε_AA_, ε_BB_, and ε_AB_) were varied in the range 0.00-1.40 k_B_T. To infer from the cHi-C data the strength of each enforced loop in the wt conditions, we assigned a rank r*_i_* (from 0 to N_loops_-1) to each detected loops (see Section **Analysis of looping interactions**) by ordering them for decreasing number of total counts (we consider as loop anchors 10 kb-regions around the central nucleotide of the detected looping region). The attraction strength of the loop *i*, was defined as ε_L,i_=(N_loops_- r_i_)/N_loops_ε_L_, where ε_L_ was varied in the range 0.00-4.00 k_B_T.

The loop-extrusion process was regulated by two players (**Fig. 6a**): the extruders and the barriers. In our model, extruders bind on chromatin, start two-sided extrusion, and keep extruding until the end of their lifetime or until they reach the first or the last monomer in the chain. In any of these three cases, extruders are relocated to a random position on the polymer chain. Extruders can cross each other upon encounter and the extrusion speed was set to 1kb/s (**Fig. 6a**)^36,37,95^. We tested various values of extruders’ density on chromatin, N_e_=4, 8, 16, and 32 extruders/Mb, and constant nominal extruders’ lifetime, L_e_=5-24 min. Notably, due to the finite size of the chain and the effect of barriers, the effective lifetime (the time going from the binding to the unbinding of one extruder from the model chromatin in the simulation) can differ from the nominal one. Hence, we refer and report the effective average lifetime of the simulation. The positions of loop-extrusion barriers have been determined by looking at the maxima of the insulation score (IS) profiles. The barriers could stop extruders from both directions with a probability *(1-p)*, where *p* is the permeability of the barrier. As previously proposed, once one extruder engaged with a side of an active barrier, it was paused there for a pausing time t_p_, that is barrier and direction specific, t_p_^L^ (for extruders arriving from the left side of the barrier) and t_p_^R^ (for extruders arriving from the right side of the barrier)^35^. During this pausing, the extruder could continue extruding in the unaffected direction (one-sided extrusion) until the extruder dissociates and re-associates elsewhere. Per each barrier, we tested several values of permeability in the range [0.00-1.00] and pausing time in the range [0-6 min]. The models’ particles assigned to A/B compartments, loop anchors, and loop-extrusion barriers in each condition are reported in **Supplementary Table 5**. All the set of parameters tested are reported in **Supplementary Table 6** for each of the simulated conditions.

###### Model parameterization

To test each of the parameters’ sets, we performed MD simulations for the time needed to extrude ∼500 Mb of chromatin in 50 independent replicates (**Supplementary Video 1**). Next, we computed the number of contacts per polymer bead of 5 kb within a distance threshold of 100 nm during the first half and the complete trajectories. From the models’ contact maps, we computed the number of contacts as a function of the genomic separation (P(s)), the normalized insulation score profile, and the number of contacts between the looping anchors. Finally, to compare cHi-C data and the models in each of the tested parameter sets, we applied the following metrics:

- The models’ contact maps and ICE-normalized cHi-C matrices were compared element by element using the Spearman correlation coefficient (SCC), where -1 corresponds to anti-correlation, 0 to no-correlation, and 1 to correlation.
- The models and cHi-C *cis*-decay curves (average number of contacts *vs.* genomic distance) were normalized such that P(s=50kb)=1 and compared using the Euclidean distance for genomic separations larger than 50 kb, d_P_. Finally, the Euclidean distance was normalized to the distance of the cHi-C profile with a profile always equal to zero d^0^_P_: D_Ps_=(d^0^_P_-d_P_)/d^0^_P_. The closer D_Ps_ is to 1 the better the models recover the cHi-C data.
- The models and cHi-C normalized IS profiles were compared using the Euclidean distance considering only the sites of the imposed loop-extrusion barriers +/- 5 kb, d_IS_. Finally, the Euclidean distance was normalized between 0 and 1 using the number of bins, N_bins_: D_IS_=(N_bins_-d_IS_)/N_bins_. The closer D_IS_ is to 1 (0) the better (worse) the models recover the cHi-C data.
- The models and cHi-C ICE-normalized maps were converted into maps of the ranks of each entry (the largest entry has rank NxN and the lowest entry has rank 1). Entries with the same value were associated with the lowest possible rank. Next, we looked at the ranks of the site-specific looping interactions and compared the distributions of the ranks of models and cHi-C looping interactions using a Wilcoxon statistical test (H0: true median shift is equal to zero. The two variables are not normally distributed). In this case, we posed that the distribution computed on the models shouldn’t be statistically different from the cHi-C one, p*-value* (p_L_)>0.10.

Given the considerable number of parameters to optimize we decided to follow a supervised optimization strategy. Starting with the ESC and NPC wt conditions, we tested the compartment-driven attractions alone. These preliminary tests showed that the simulations with purely homotypic interactions (ε_AB_=0) or not-equal *cis-* and *trans-*attractions compartment attractions (ε_AA_=ε_BB_ and ε_AA_=ε_AB_) don’t recover the cHi- C P(s) **Extended Data Fig. 7a-b**. Next, we tested the loop-extrusion process varying N_e_ and L_e_ with ε_AA_=ε_BB_=ε_AB_=0 with no barriers and found constraints for N_e_<8 and L_e_∼20min (**Extended Data** Fig. 7c). We, then, optimized the permeabilities and pausing-time at barriers with a supervised approach first varying the parameters for the barriers with higher insulation scores, B^R^ and B^L^, that delimit the large central TAD in the region and then for all the others (**Extended Data Fig. 7d-e**). Finally, we optimize the strength of the site-specific looping attractions, ε_L_. This procedure resulted in a few sets of optimized parameters per condition, which differ for small adjustment of two or three parameters and resulted in very similar values of the optimized metrics. In the following, we list the optimized parameters’ sets with the correspondent range of values of the four metrics:

<u>ESC wt</u> : ε_AA_=ε_BB_=ε_AB_=0.00-0.02 k_B_T, ε_L_=3.00 k_B_T, N_e_=8 Extruders/Mb, L_e_=9min, p=(0.95, 0.50, 0.75, 0.70, 0.30, 0.95), t_p_^L^=(30, 30, 0, 0, 60, 30), t_p_^R^=(30, 300, 0, 0, 30, 30), T_sim_=160 and 320 min, SCC_m_=0.82-0.83, D_Ps_=0.78-0.80, D_IS_=0.92-0.94, p_L_=0.81-0.88.

<u>NPC wt</u> : ε_AA_=ε_BB_=ε_AB_=0.00-0.02 k_B_T, ε_L_=3.00 k_B_T, N_e_=8 Extruders/Mb, L_e_=12min, p=(0.85, 0.30, 0.80, 0.75, 0.85, 0.30, 0.90), t_p_^L^=(30, 30, 0, 0, 30, 60, 30), t_p_^R^=(30, 300, 0, 0, 30, 30, 30), T_sim_=160 and 320 min, SCC_m_=0.82-0.84, D_Ps_=0.82-0.84, D_IS_=0.90-0.96, p_L_=0.90-0.93 and 0.74.

To model the NPC mutations, we perturbed the optimal parameters found in NPC wt. For the NPC Δprom and NPC polyA conditions, we tested an increase of the permeabilities at 3’ and promoter by 0.05, 0.10 and 0.15, an increase of looping strength ε_L_=4 k_B_T, and a tuning down of the promoter-anchored looping attractions. For NPC ΔA, we set the permeability and lifetimes at A, and strength of A-anchored looping interactions to 0 and tested a tuning-down of the promoter-anchored looping interactions. For NPC ΔAB, in addition to the NPC ΔA perturbations, we also set the strength of B-anchored looping interactions to 0 and tuned the permeability at the promoter to 0.65 to account for the increased insulation at this point. For NPC ΔB, we set the strength of B-anchored looping interactions to 0 and tested a decrease of the permeability at the promoter and of the promoter-anchored looping interactions. For ΔCTCF_A_, we tested an increase of the permeability and a decrease of the looping interactions at A, and a decrease of the permeability at the promoter and 3’. To tune-down the looping attractions anchored at the Promoter or A we introduced a weighting factor (L_Pf_ and L_Af_ respectively) of 0.25, 0.50, or 0.75 to the attraction strength value. In the following, we list the optimized parameters’ sets with the corresponding rangess of the four metrics. In bold, we show the parameters that are different from NPC wt condition (**Extended Data** Fig. 8).

<u>NPC Δprom</u> : ε_AA_=ε_BB_=ε_AB_=0.00k_B_T, **ε_L_=4.00k_B_T**, **L_Pf_=0.75**, N_e_=8Extruders/Mb, L_e_=13min, p=(0.85, 0.30, **0.85**, **0.85**, 0.85, 0.30, 0.90), t_p_^L^=(30, 30, 0, 0, 30, 60, 30), t_p_^R^=(30, 300, 0, 0, 30, 30, 30), T_sim_=320min, SCC_m_=0.91, D_Ps_=0.81-0.83, D_IS_=0.94, p_L_=0.49-0.58.

<u>NPC polyA</u> : ε_AA_=ε_BB_=ε_AB_=0.02k_B_T, ε_L_=3.00k_B_T, **L_Pf_=0.25**, N_e_=8Extruders/Mb, L_e_=14min, p=(0.85, 0.30, **0.85**, **0.85**, **0.90**, 0.30, 0.90), t_p_^L^=(30, 30, 0, 0, 30, 60, 30), t_p_^R^=(30, 300, 0, 0, 30, 30, 30), T_sim_=160-320min, SCC_m_=0.94, D_Ps_=0.86-0.87, D_IS_=0.93, p_L_=0.74.

<u>NPC ΔA</u> : ε_AA_=ε_BB_=ε_AB_=0.02k_B_T, ε_L_=3.00k_B_T, **L_Pf_=0.75**, N_e_=8Extruders/Mb, L_e_=13min, p=(0.85, 0.30, 0.80, 0.75, **1.00**, 0.30, 0.90), t_p_^L^=(30, 30, 0, 0, **0**, 60, 30), t_p_^R^=(30, 300, 0, 0, **0**, 30, 30), T_sim_=160-320min, SCC_m_=0.95, D_Ps_=0.85-0.86, D_IS_=0.90-0.91, p_L_=0.13-28.

<u>NPC ΔAB</u> : ε_AA_=ε_BB_=ε_AB_=0.02k_B_T, ε_L_=3.00k_B_T, **L_Pf_=0.75**, N_e_=8Extruders/Mb, L_e_=13min, p=(0.85, 0.30, 0.80, **0.65**, **1.00**, 0.30, 0.90), t_p_^L^=(30, 30, 0, 0, **0**, 60, 30), t_p_^R^=(30, 300, 0, 0, **0**, 30, 30), T_sim_=160-320min, SCC_m_=0.94, D_Ps_=0.81-0.83, D_IS_=0.93-0.94, p_L_>0.09-0.15.

<u>NPC ΔB</u> : ε_AA_=ε_BB_=ε_AB_=0.02k_B_T, ε_L_=3.00k_B_T, **L_Pf_=0.75**, N_e_=8Extruders/Mb, L_e_=13min, p=(0.85, 0.30, 0.80, **0.70**, 0.85, 0.30, 0.90), t_p_^L^=(30, 30, 0, 0, 30, 60, 30), t_p_^R^=(30, 300, 0, 0, 30, 30, 30), T_sim_=160-320min, SCC_m_=0.82, D_Ps_=0.84-0.85, D_IS_=0.95, p_L_>0.15-0.19.

<u>NPC ΔCTCF_A_</u> : ε_AA_=ε_BB_=ε_AB_=0.00k_B_T, ε_L_=3.00k_B_T, **L_Af_=0.50**, N_e_=8Extruders/Mb, L_e_=13min, p=(0.85, 0.30, 0.80, 0.75, **0.90**, 0.30, 0.90), t_p_^L^=(30, 30, 0, 0, 30, 60, 30), t_p_^R^=(30, 300, 0, 0, 30, 30, 30), T_sim_=160-320min, SCC_m_=0.86, D_Ps_=0.86-0.87, D_IS_=0.91, p_L_=0.46-0.71.

We notice that the gain in correlation respect to the the NPC wt condition may be limited in some conditions, but still quantifiable at a visual inspection of the compared quantities (**Extended Data Fig. 7e**). The parametrization procedure was conducted by testing a total of 740 parameters’ sets: 336 for ESC wt, 252 for NPC wt, 36 for NPC Δprom, 20 for NPC polyA, 16 for NPC ΔA, 12 for NPC ΔAB, 24 for NPC ΔB, and 44 for NPC ΔCTCF.

## Acknowledgements

We thank Peter Hansen for his help with cHi-C design. We thank Pia Mach, Pavel Kos, and Daniel Jost for fruitful discussions on biophysical modelling. We are grateful to the Genotoul bioinformatics platform Toulouse Occitanie (Bioinfo Genotoul, https://doi.org/10.15454/1.5572369328961167E12) and the Pôle Scientifique de Modélisation Numérique (PSMN) of the ENS de Lyon for computational resources. We thank the Montpellier Ressources Imagerie facility (BioCampus Montpellier, Centre National de la Recherche Scientifique (CNRS), INSERM, University of Montpellier) for their help and support with microscopy. IJ was supported by EMBO Long-term Fellowship ATLF 559-2018. MDS was supported by an advanced ERC grant “3DEpi” and an ANR “live chrome”. HR was supported by La Ligue Contre le Cancer. MS was supported by a Marie Skłodowska-Curie Innovative Training Network (grant no. 813327 ‘ChromDesign’) and Fondation de Recherche Medicale. D.N., F.B., G.L.P. and G.C. were supported by the CNRS. The laboratory of G.C. was supported by grants from the European Research Council (Advanced Grant 3DEpi, under grant agreement number 788972), the European Union (CHROMDESIGN Project, under the Marie Skłodowska- Curie grant agreement number 813327), the Fondation pour la Recherche Médicale (EQU202303016280), the MSDAVENIR foundation (project GENE- IGH), the INSERM, the Centre National pour la Recherche Scientifique, the Agence Nationale de la Recherche (E- RARE project ‘IMPACT’ and “PLASMADIFF3D” project, ANR-18-CE15-0010) and by the French National Cancer Institute (INCa PLBIO18-362).

## 7. Contributions

GC and IJ conceived the study. IJ designed and carried out wet-lab experiments. IJ analysed ChIP-seq and RNA-seq data. MDS designed and carried out biophysical modelling. MDS and GPL analysed the cHi-C data. MS helped carry out ChIP-seq and with the RNA-seq analysis. HR and FB designed and together with DN carried out 3D FISH. HR and FB analysed the 3D-FISH. IJ and MDS wrote the manuscript with the input of all other authors.

## 8. Competing Interests

The authors declare no competing interests.

## 9. Data Availability

All raw data were submitted to the National Library of Medicine’s (NCBI) Sequence Read Archive (SRA) and the processed files were submitted to the Gene Expression Omnibus (GEO). All data can be retrieved under the GEO SuperSeries number **GSE262551**. RNA-seq processed data (FPKMs and differentially expressed genes) can be found in the Supplementary Tables 1 and 2 and counts for loop quantification can be found in Supplementary Table 8.

## 10. Code Availability

Scripts used in this article will be made available at Cavalli lab GITHUB page.

---

**Extended Data Figure 1.**
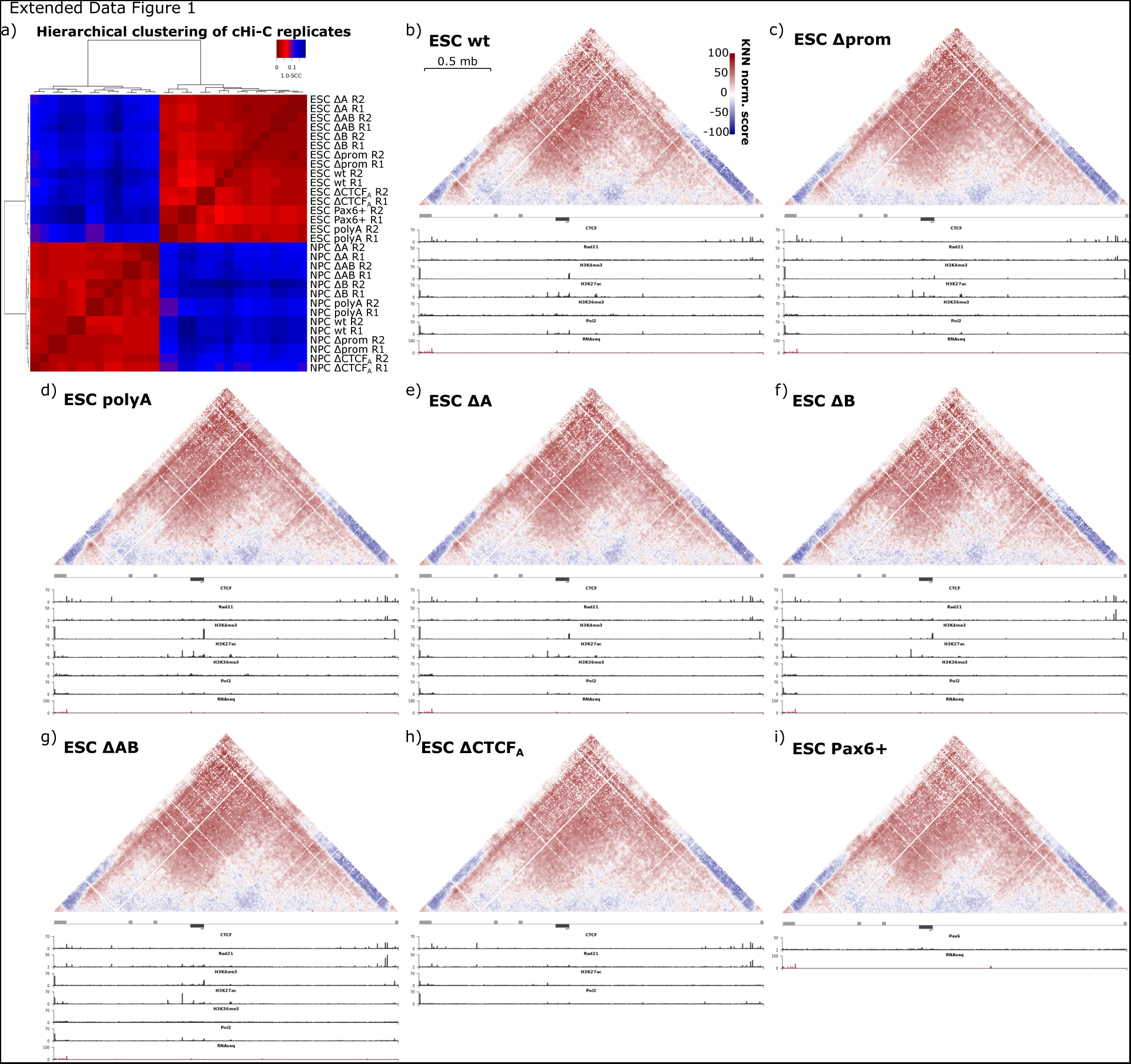
ESC cHi-C maps, ChIP-seq and RNA-seq profiles in wt and mutants. **a)** Hierarchical clustering analysis of the <u>S</u>tratum-adjusted <u>C</u>orrelation <u>C</u>oefficient (SCC) from the *HiCRep* package to compare the cHi-C biological replicates (**Methods**). **b-i)** cHi-C score (observed/expected) maps with ChIP-seq data (CTCF, RAD21, H3K4me3, H3K27Ac, H3K36me3, Pol2) and RNA-seq data depicted below the maps for **b)** ESC wt, **c)** ESC Δprom, **d)** ESC polyA, **e)** ESC ΔA, **f)** ESC ΔB, **g)** ESC ΔAB, **h)** ESC ΔCTCF_A_, and **i)** ESC Pax6+ conditions at 5 kb resolution. For e) shown ChIP-seq tacks are: CTCF, RAD21, H3K27Ac, and Pol2 and no RNA-seq. For h) only PAX6 ChIP-seq and RNA-seq are present.

**Extended Data Figure 2.**
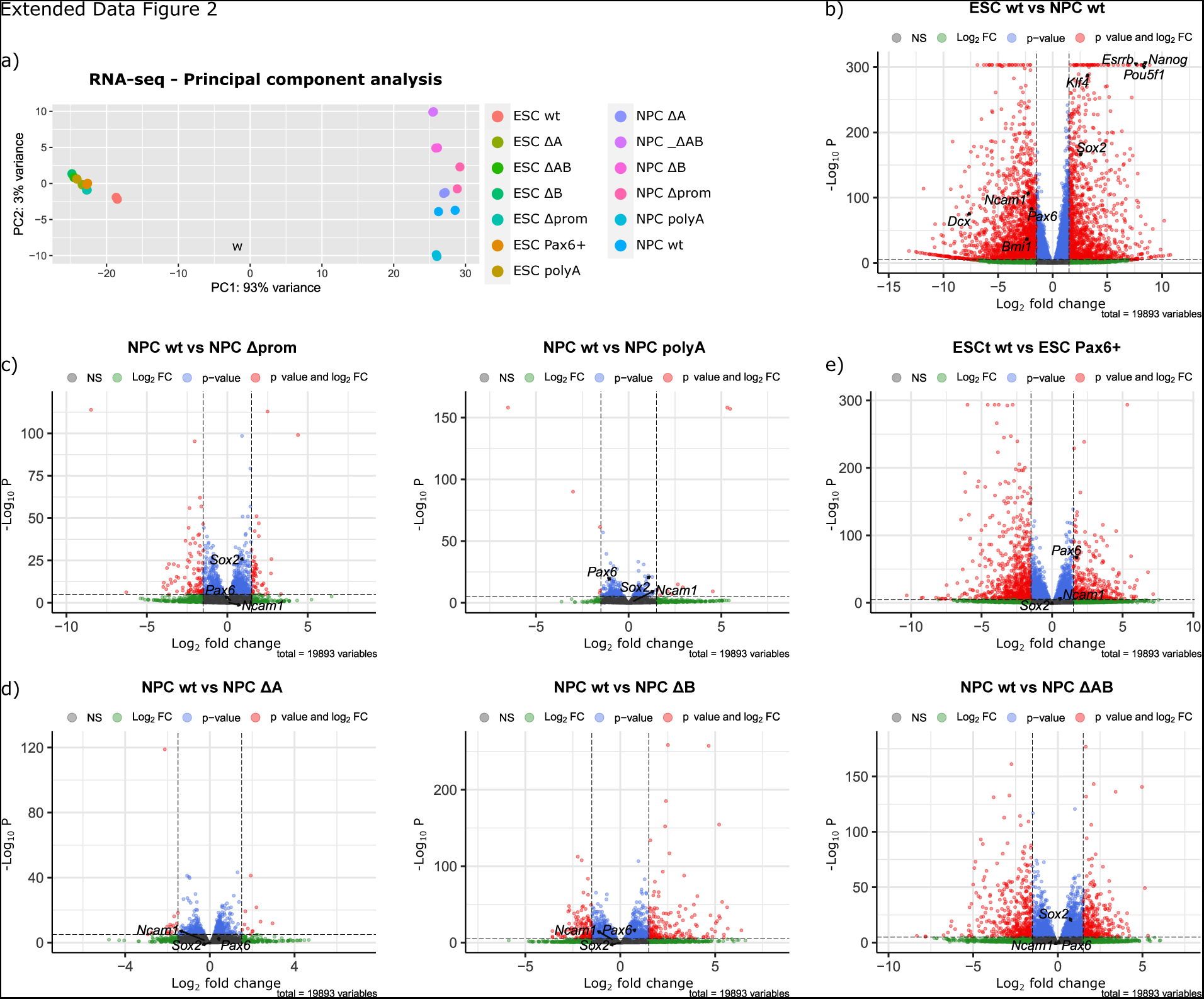
Total RNA-seq expression changes. **a)** Principal component analysis (PCA) of all the RNA-seq biological replicates in all the examined conditions. **b)** DEseq2 pairwise analysis of ESC wt and NPC wt conditions with depicted main pluripotency factors (*Pou5f1* (*Oct4*), *Esrrb*, *Nanog*, *Klf4*, *Sox2*) and NPC and neuronal lineage markers (*Pax6*, *Dcx*, *Bmi1*, *Ncam1*). **c)** DEseq2 pairwise analysis of NPC wt and either NPC ΔA, ΔB or ΔAB conditions with depicted main NPC markers (Pax6, *Sox2*) and NSC marker (*Ncam1*) to indicate that all conditions remain of the same NPC identity. **d)** DEseq2 pairwise analysis of NPC wt and either NPC Δprom or polyA conditions with depicted main NPC markers (*Pax6*, *Sox2*) and NSC marker (*Ncam1*) to indicate that all conditions remain of the same NPC identity. **e)** DEseq2 pairwise analysis of ESC wt and ESC Pax6+ conditions with depicted main NPC markers (*Pax6*, *Sox2*) and NSC marker (*Ncam1*) to indicate that except for the *Pax6* expression, the neuronal markers stay low and *Sox2* expression does not decrease. For detailed neuronal and pluripotent marker depiction see **Extended Data** Figure 9b.

**Extended Data Figure 3.**
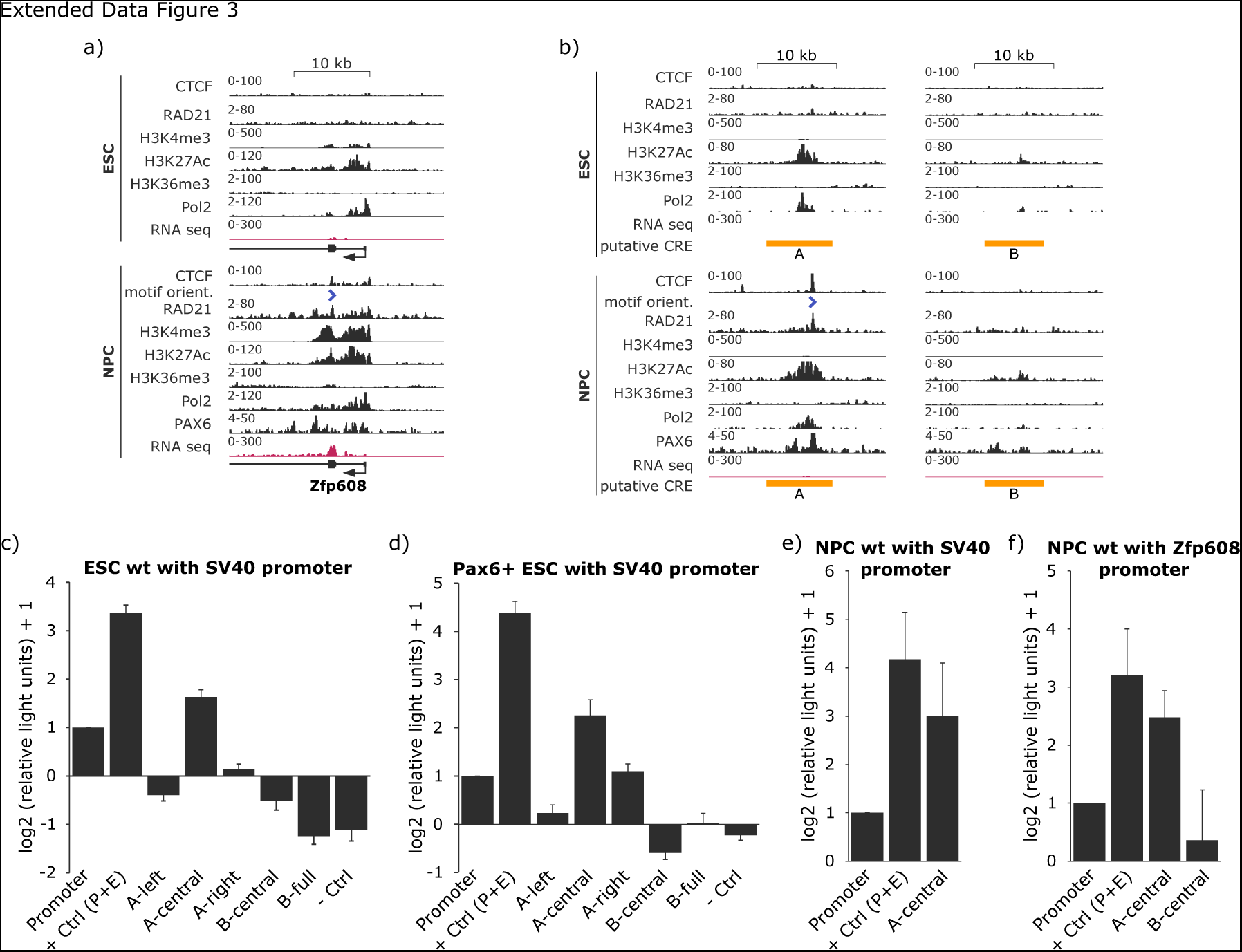
TF and histone mark presence at the *Zfp608* promoter and two putative enhancers, and luciferase assay of putative enhancers. Zoom into the TF binding and histone mark presence in ESC and NPC state at: **a)** *Zfp608* promoter, and **b)** two loop anchors (putative enhancer elements). For luciferase, element A (8.2 kb) was divided into three regions, left (A-left), central (A-central - containing all histone marks and TF binding), and right (A-right). B element was cloned either in full (B-full - 7.4 kb) or only the central region (B-central that carries the H3K27Ac signal). +Ctrl (P+E) is a construct carrying SV40 promoter and SV40 enhancer. -Ctrl is a construct carrying a random region occupied by H3K9me3. **c)** Luciferase assay in ESC wt cells using constructs with SV40 promoter. **d)** Luciferase assay in ESC Pax6+ (ESCs ectopically expressing Pax6) cells using constructs with SV40 promoter. **e)** Luciferase assay in NPCs using constructs with SV40 promoter. **f)** Luciferase assay in NPCs using constructs with *Zfp608* cognate promoter.

**Extended Data Figure 4.**
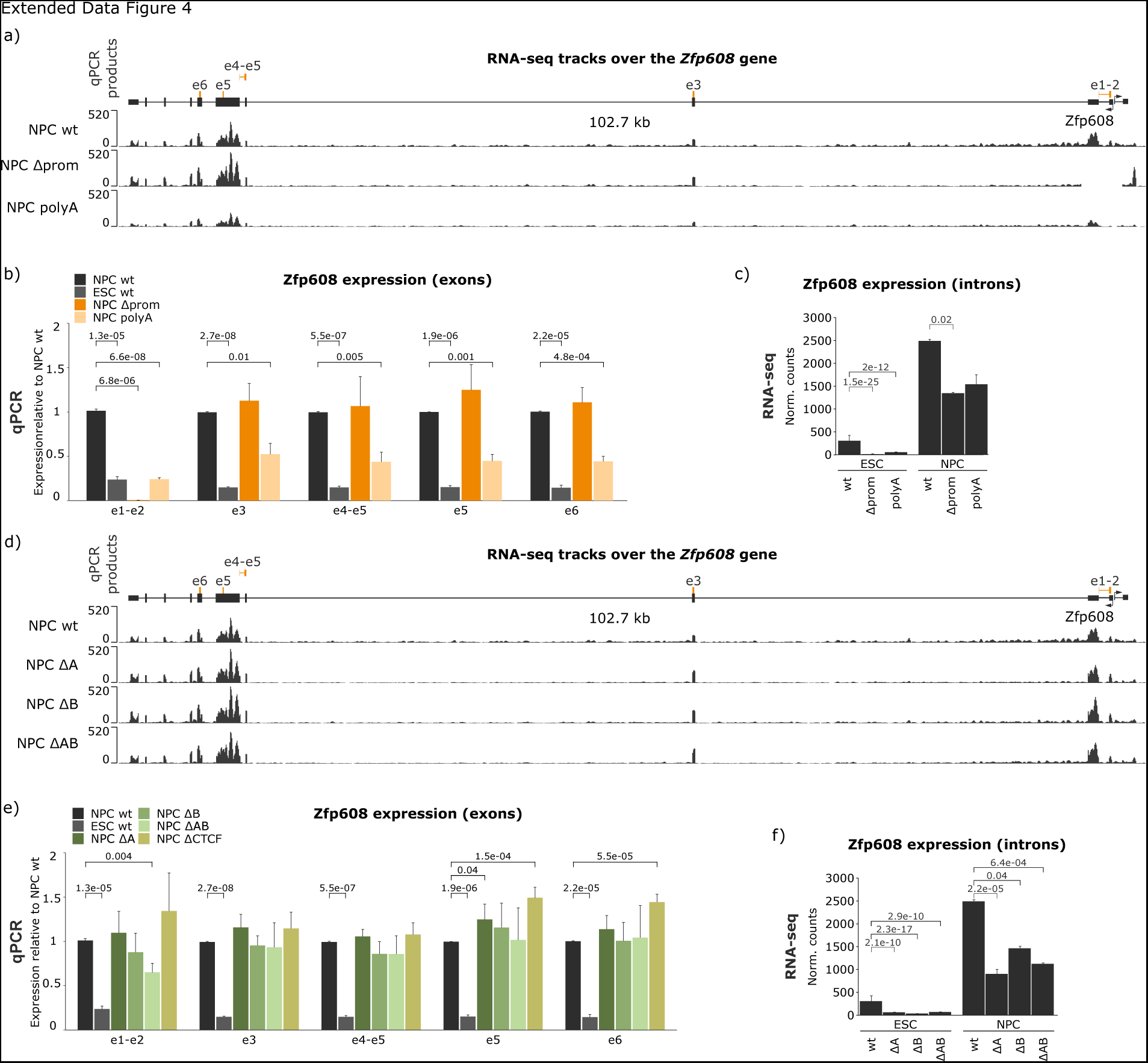
*Zfp608* expression changes in all mutants. **a)** RNA-seq profiles of NPC wt, ΔA, ΔB, and ΔAB over the *Zfp608* gene. At the top of the RNA-seq profiles, we show the detailed annotation of the *Zfp608* gene and the location of the real-time quantitative PCR (RT-qPCR) primers used in b). **b)** RT-qPCR results for *Zfp608* expression using primers from a) and using at least three biological replicates except for ΔCTCF_A_ which was done in two biological replicates. p-values were calculated using paired two-sample Student’s t-test. **c)** Deseq2 RNA-seq normalized counts over all *Zfp608* introns in ESC and NPC wt, ΔA, ΔB, and ΔAB conditions. p_adj_ values are derived from DEseq2 pairwise comparison (Wald test) of the wt with the corresponding mutant in the corresponding cell line (i.e., NPC wt vs NPC ΔA). **d)** RNA-seq profiles of NPC wt, Δprom, and polyA over the *Zfp608* gene. At the top of the RNA-seq profiles we show detailed annotation of the *Zfp608* gene and the location of the RT-qPCR primers used in e). **e)** RT-qPCR results for *Zfp608* expression using primers from d) and using two biological replicates for mutants and at least three for the NPC and ESC wt. p-values were calculated using paired two-sample Student’s t-test **f)** RNA-seq normalized counts over all *Zfp608* introns in ESC and NPC wt, Δprom, and polyA conditions. p_adj_ values are derived from DEseq2 pairwise comparison (Wald test) of the wt with the mutants in the corresponding cell line (i.e., NPC wt vs NPC Δprom).

**Extended Data Figure 5.**
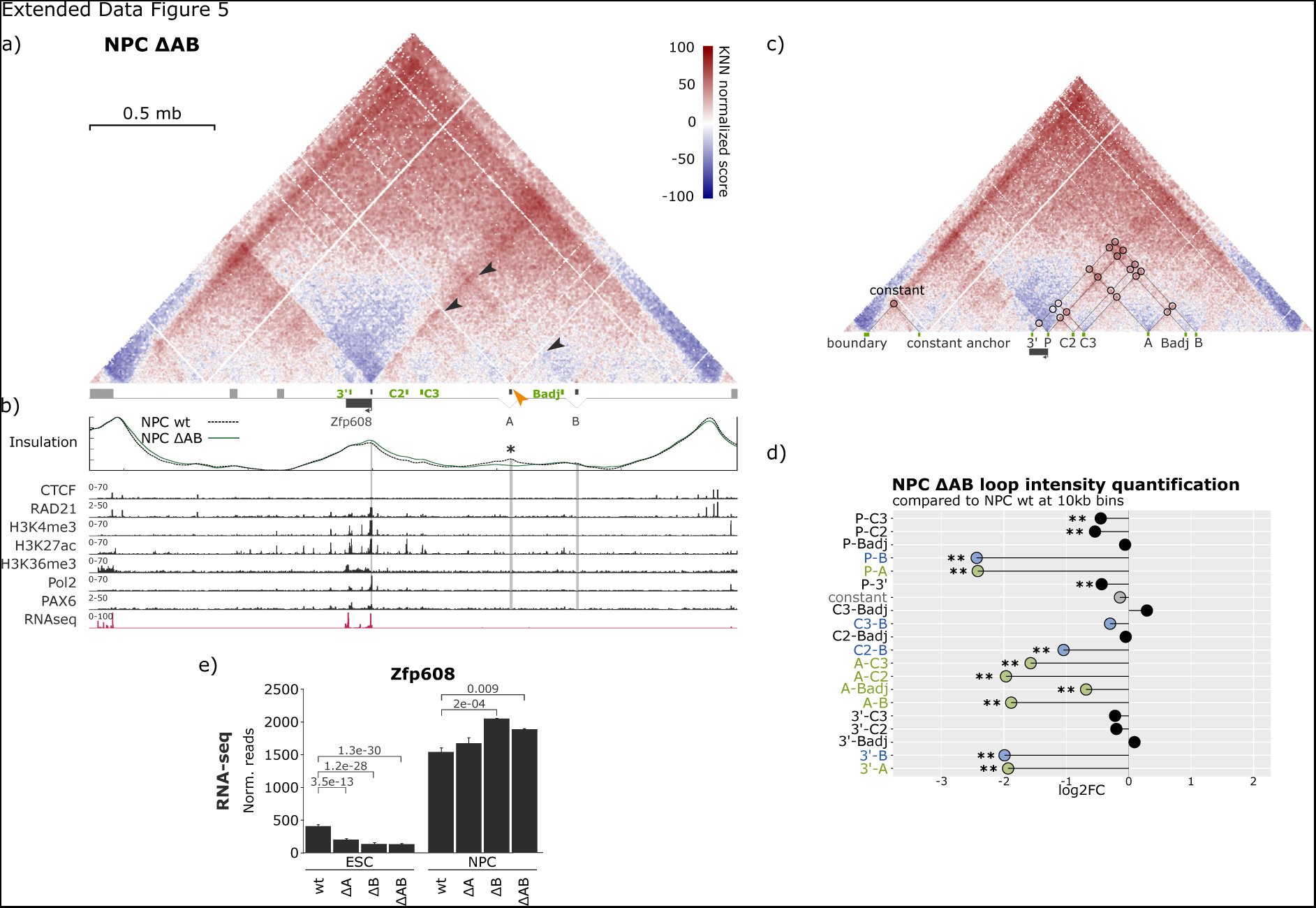
Impact of double element AB deletion on local chromatin rewiring. **a)** NPC cHi-C score maps (observed/expected) of the double element AB deletion (ΔAB; 8.2 kb and 7.4 kb deletions, respectively) at 5 kb resolution. Three NPC-specific contacts are annotated with black arrows (P-A, P-B, and A-B). Orange arrows indicate visible changes in the insulation pattern between the mutant and wt. **b)** The top panel depicts the experimentally quantified insulation score at the capture region for ΔAB. The asterisk indicates sites of significantly changed IS between the mutant and wt condition calculated by Welch Two Sample t-test; ΔAB condition at enhancer A - pval=6.5E-12. The panel below shows ChIP-seq (CTCF, RAD21, H3K4me3, H3K27Ac, H3K36me3, Pol2, and PAX6), and RNA-seq data. **c)** Circles on an NPC wt cHi-C map show the main NPC-specific putative E-P and E-E contacts. Below the map, the anchors of each contact (loop) are named so that the loop can easily be identified in panel d) (*i.e.*, P-A is a loop with one anchor in P and the other in A). **d)** log2FC of the loop intensity in NPC ΔAB mutant over the NPC wt condition (**Methods**). In green all loops that have one anchor in A are annotated and in blue all loops that have one anchor in B are annotated. * indicates loops with 0.01<*p*<0.05, and ** indicate *p*<0.01, where the *p*-values of the log2FC were calculated from a Wald statistical test with DEseq2 (**Methods**). **e)** Deseq2 normalized RNA-seq counts of ESC and NPC wt, ΔA, ΔB, and ΔAB cell lines and their p_adj_ values. p_adj_ was calculated using DEseq2 (Wald test) by pairwise comparison of individual mutants to the wt of the corresponding cell line (*i.e.*, ESC wt to ESC ΔA).

**Extended Data Figure 6.**
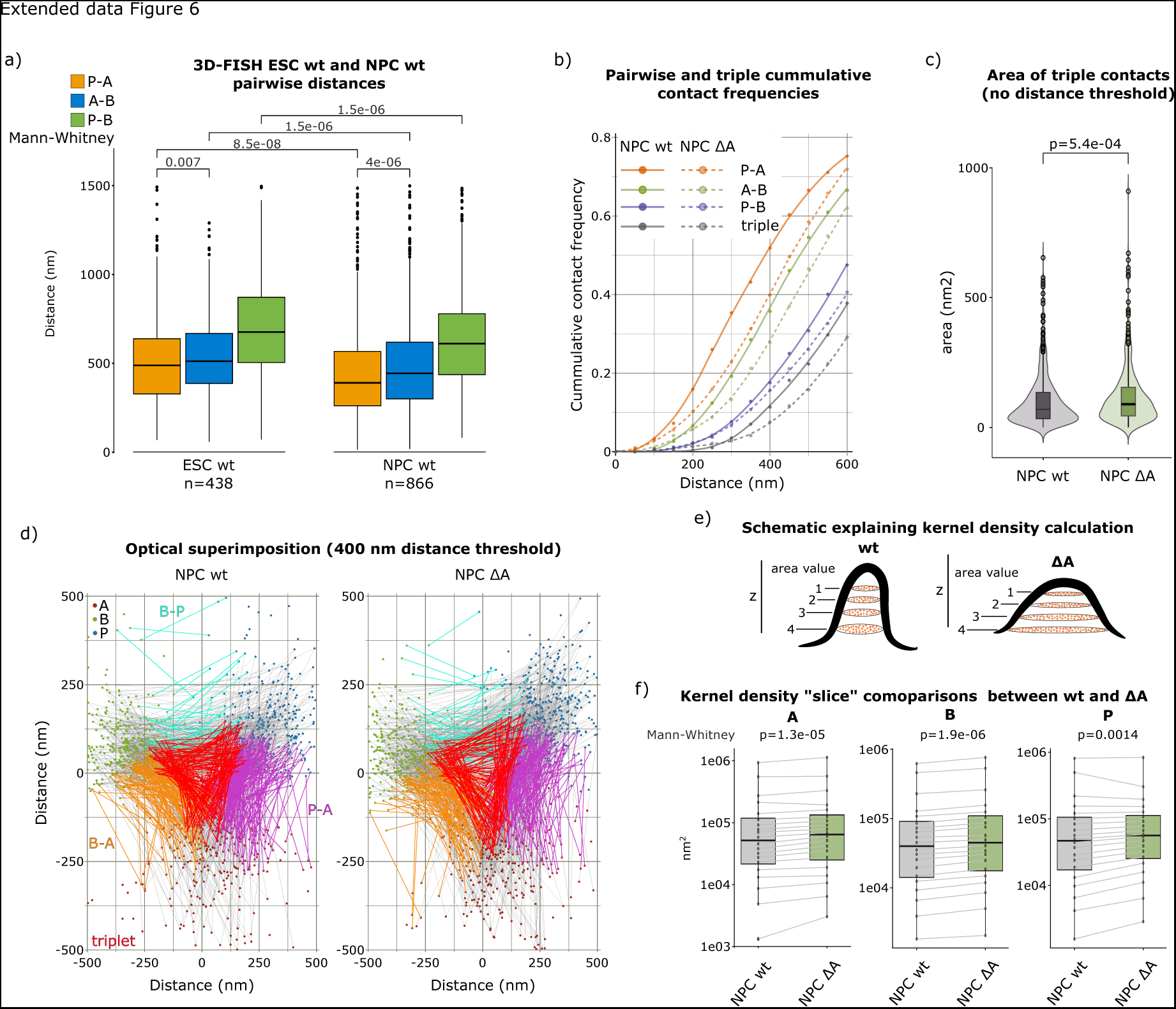
3D FISH in NPC wt, NPC ΔA, and ESC wt conditions. **a)** Pairwise (P-A, P-B, and A-B) distances in ESC wt and NPC wt. p-values (indicated above each pairwise comparison) were calculated using the Mann-Whitney statistical test. In addition, a Mann-Whitney statistical test was calculated for the distanced A-B and P-A in ESC wt and NPC wt to quantify how significant is observed P-A compaction in each state. **b)** Pairwise (P-A, P-B, and A-B) and triple-contact cumulative frequencies (P-A-B present in the same cell) as a function of the distance thresholds **(Methods)**. P-A is depicted in orange, A-B in green, P-B in blue, and triple-contacts in grey. The bold line represents NPC wt and the dashed line NPC ΔA. Triple-contacts are determined if each of the three pairwise distances is under a certain threshold (i.e., if P-A, A-B, and P-B distances in the same cell are all under 400 nm then it is considered a triple-contact at that threshold). This plot is an extension of Fig. 4c showing all pairwise-distance curves. **c)** Area of the triangle formed by P, A, and B calculated with no distance threshold for pairwise contacts (all distances) **(Methods)**. p-value was calculated using the Mann-Whitney statistical test. This is an extension of Fig. 4d, which was obtained imposing a distance threshold of 400 nm. **d)** Position of P, A, and B genetic elements optimally superimposed on a plane. Briefly, distances between elements were measured and conserved while reporting them on a single plane by aligning the barycenter of every triangle. In addition, every genetic element was projected at the same side so that element B is always top left, promoter top right, and element A bottom center. The elements are also labeled by colour, green dots: element B, red dots: element A, and the blue dots: promoter (P). Cyan lines connect all P-B pairs that are under 400 nm, purple lines connect all P-A pairs that are under 400 nm, and orange lines connect all A-B pairs under 400 nm. Red triangles connect P, A, and B elements if each of the pairwise distances is under 400 nm, thus representing triple contacts at this distance threshold. The area of these red “triangles” is shown in Fig. 4d. **e)** A schematic drawing of how the kernel density map was generated. Briefly, the positions of genetic elements and their preserved distances were plotted on a plane as explained in d). The kernel of a 2D distribution is a set of values defined on each pixel of the plane. The sum of these values over the entire plane equals 1. If we consider a single pixel, the value of the kernel is equal to the value of that pixel divided by the area of the pixel and by the total number of points. Since the area of the pixel and the total number of points are the same for each of the 3 elements (P, A, and B), we can compare the values of the 2D kernel between the three distributions. For this, every element was divided in increments of 5% on the z-axis (20 stacks) so that for every z-stack (labeled 1-4 on the schematic) area could calculated and compared directly for the three elements, represented in f). **f)** The area values of the same stack were connected with lines so that the change in area value could be seen and compared for each of the 20 z-stacks. Then, the Mann-Whitney paired test with Bonferroni correction for multiple testing was used to calculate a p-value of these changes between NPC wt and NPC ΔA conditions.

**Extended Data Figure 7.**
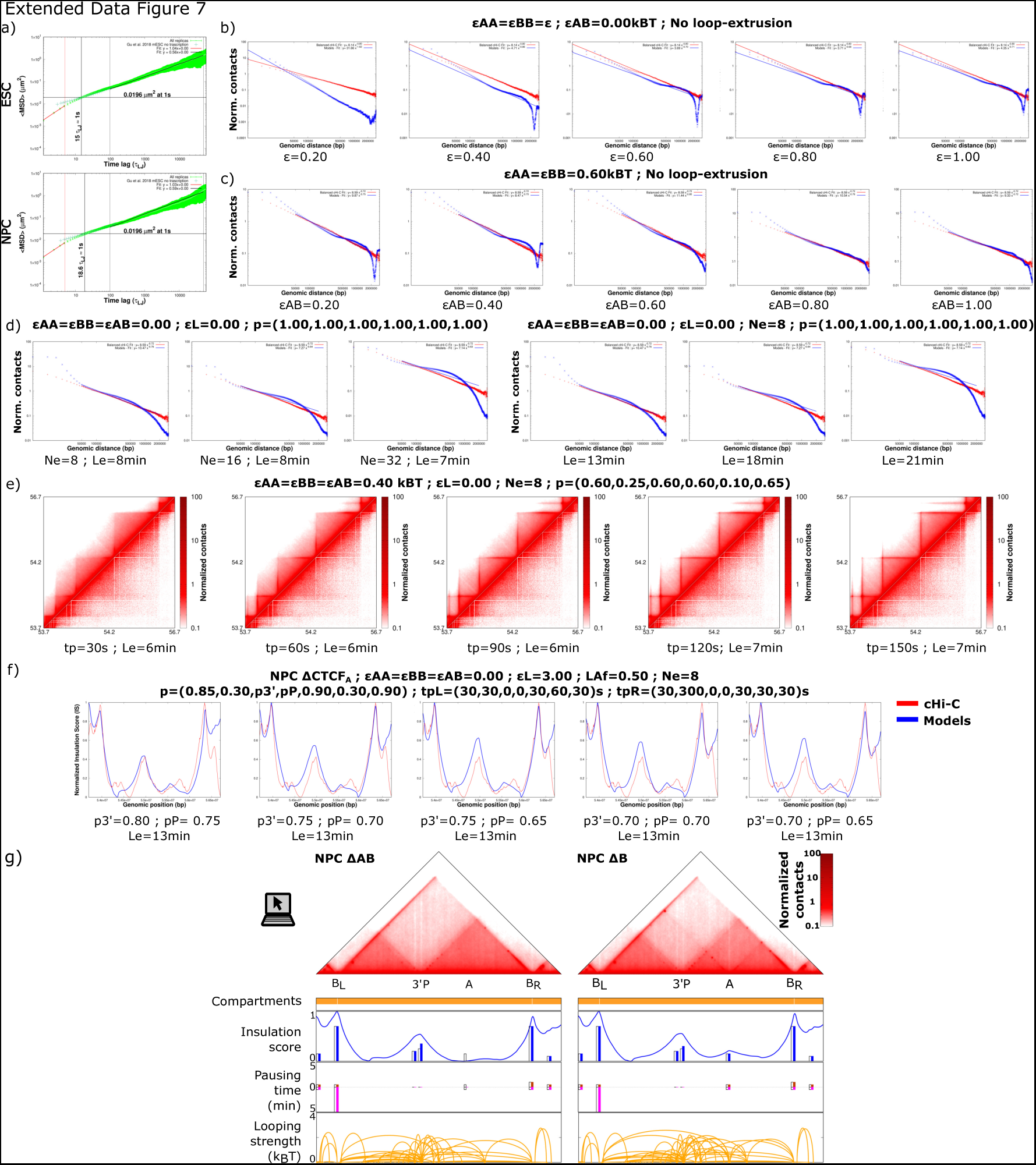
Summary of selected sets of parameters used in the biophysical modelling. Selection of parameters sets explored in the optimization of the biophysical models. **a)** Comparison of the mean-squared displacement (MSD) as a function of simulation time in the ESC and NPC systems with experimental data from Gu et al. for non-transcribed genes^1^. **b)** *cis*-decay curves for ESC wt modeling (*blue*) obtained using increasing compartment-specific (E_AB_=0) attraction strengths show discrepancies with the cHi-C cis-decay curves around a genomic distance of 2.4 Mb. **c)** *cis*-decay curves for ESC wt modeling (*blue*) obtained using compartment self-attraction strength of E_AA_=E_BB_=0.60 k_B_T and increasing trans-compartment (E_AB_) attraction show that the discrepancies with the cHi-C *cis*-decay curves are mitigated if E_AB_ gets closer to the compartment self-attraction strength. **d)** *cis*-decay curves for ESC wt modeling obtained using increasing extruder density (*left*) and extruder lifetime (*right*) show discrepancies with the cHi-C *cis*-decay curves at genomic distance between 100 kb and 1 Mb. These tests were done in the absence of compartment attraction, loop-extrusion barriers, and site-specific looping attraction. **e)** Selection of models’ (*top left triangle*) and ICE-normalized (*bottom right triangle*) maps of the contacts from parameters’ sets with increasing pausing times at barriers in NPC wt. These plots show that pausing times larger than 30 s at all barriers generate too strong flares and barrier-barrier loops in the models’ maps. Contacts are normalized by the average number of contacts at a genomic distance of 50 kb. **f)** Normalized insulation profiles leading to the optimization of the permeabilities at the loop-extrusion barriers located at the 3’ and the promoter of the *Zfp608* gene illustrate that a variation of only 5% in the permeability values at individual barriers generates visibly distinct insulation profiles. **g)** The maps of normalized contact ranks at 5kb resolution obtained from the optimal biophysical models generated for NPC ΔAB and ΔB are shown on the top (**Methods**). The four graphs below the maps show the features of the experimental maps used as input for the simulations together with the parameters quantified using biophysical modeling. These include: the top panel) the compartments partition inferred from the Hi-C dataset from Bonev et al.; the second panel) the position of the loop-extrusion barriers marked as bars whose heights corresponds to the optimized permeabilities together with the insulation score profile obtained from the models’ map; the third panel) the left (*positive values*) and right (*negative values*) pausing times at each of the loop-extrusion barriers; and the fourth panel) loops detected by experimental cHi-C maps (**Methods**) are shown as arcs whose height indicates the corresponding optimized attraction strength. The barplots shown in colors are the permeablities and pausing times for the mutants and in white the corresponding quantities in the NPC wt condition.

**Extended Data Figure 8.**
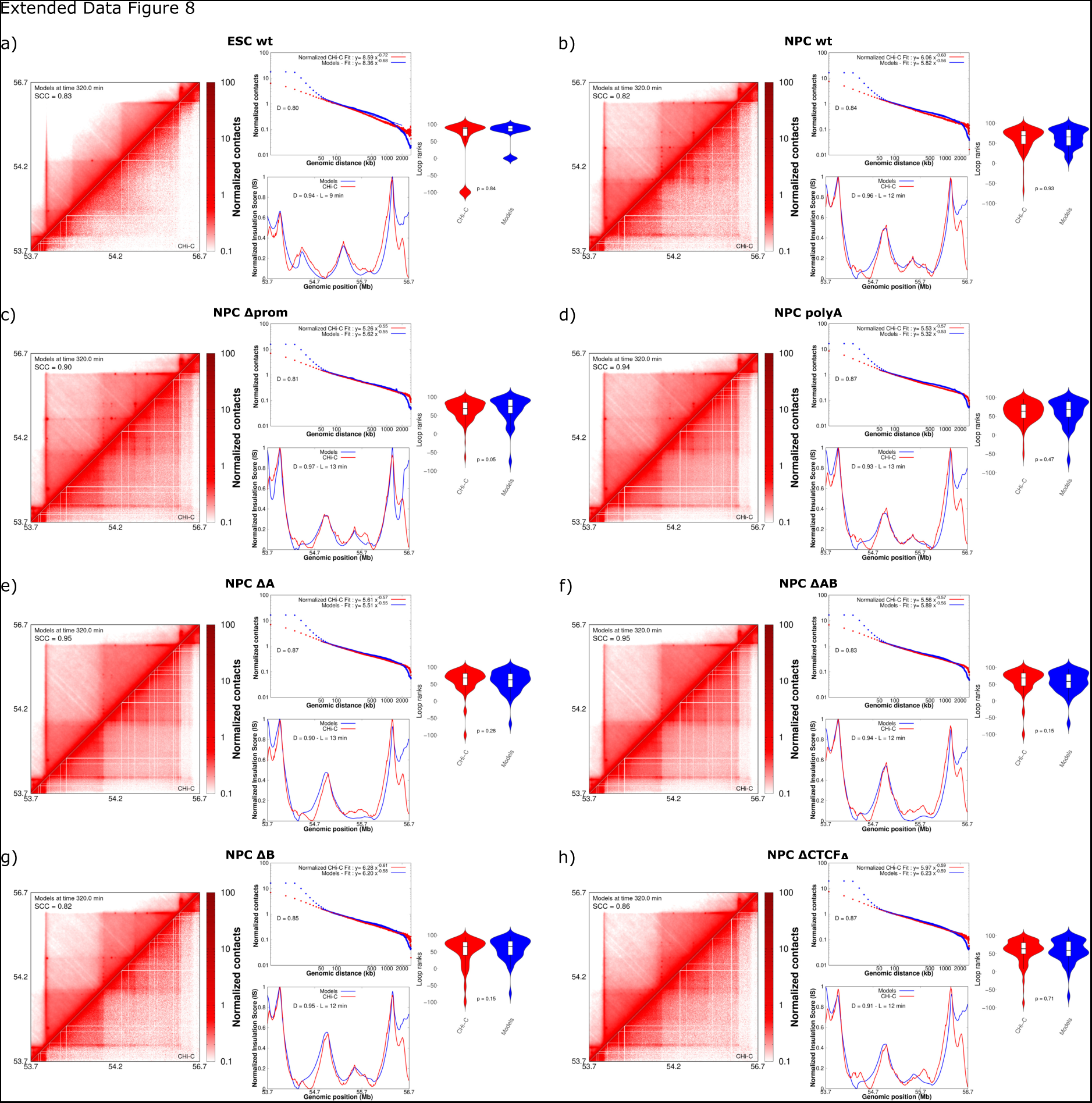
Optimized parameters for biophysical modelling of every mutant. Illustration of the metrics resulting from the comparison of the optimized biophysical models and the cHi-C data for **a)** ESC wt, **b)** NPC wt, **c)** Δprom, **d)** polyA, **e)** ΔA, **f)** ΔAB, **g)** ΔB, **h)** ΔCTCF_A_. Each of the panels shows quantities from the models’ conformations and the cHi-C data in the region chr18:53,700,000-56,700,000 bp: (*left*) the models’ (*top left triangle*) and ICE-normalized (*bottom right triangle*) maps of the contacts that are normalized by the average number of contacts at a genomic distance of 50 kb (**Methods**), (*center top*) the cis-decay curves (average number of contacts *vs.* genomic distance) computed from the models’ (*blue*) and cHi-C contact maps (*red*), (*centre bottom*) the normalized IS profiles from the models’ (*blue*) and cHi-C contact maps (*red*); (*right*) the distribution of the contact ranks corresponding to the target loops in the ESC wt (panel a) or NPC wt (panels b-h) conditions (**Methods**).

**Extended Data Figure 9.**
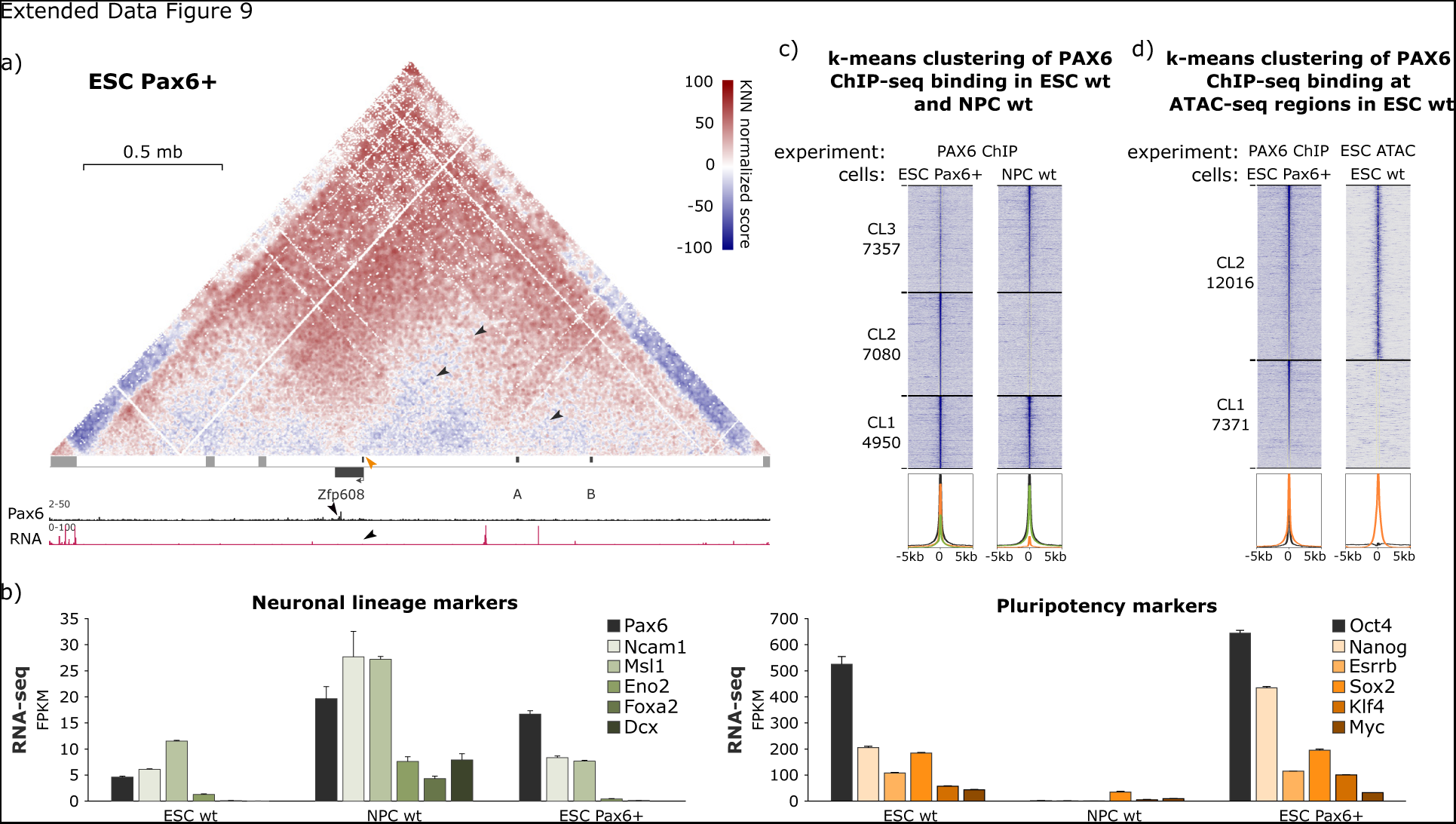
Pax6 overexpression in ESCs. **a)** cHi-C score (observed/expected) maps of the *Pax6* overexpression at the ESC state at 5 kb resolution. The expected positions of three NPC-specific contacts are annotated with black arrows (P-A, P-B, and A-B). The orange arrow annotates the position of the NPC-specific point of insulation at the *Zfp608* promoter. Below the cHi-C map, the first panel shows the ChIP-seq profile of PAX6 binding at the *Zfp608* locus and the second panel shows the RNA-seq profile at the *Zfp608* locus in the ESC Pax6+ cells. **b)** FPKM values from RNA-seq data for the main neuronal lineage and pluripotency markers. **c)** k-means clustering of global PAX6 binding in ESC Pax6+ and NPC wt cells. **d)** k-means clustering of PAX6 binding sites in ESC Pax6+ cells compared to the ESC wt open regions.

## SUPPLEMENTARY VIDEOS CAPTIONS

**Supplementary Video 1**. Illustration of the metrics resulting from the comparison of the optimized biophysical models and the cHi-C data for NPC wt over the simulation time. Each of the panels shows quantities from the models’ conformations and the cHi-C data in the region chr18:53,700,000-56,700,000 bp. From left to right we show: an example (replica 1) of the 50 molecular dynamics trajectories for the optimal NPC wt parameters’ set; the models’ (*top left triangle*) and ICE-normalized (*bottom right triangle*) maps of the contacts that are normalized by the average number of contacts at a genomic distance of 50 kb (**Methods**); (*top*) the *cis*-decay curves (average number of contacts *vs.* genomic distance) computed from the models’ (*blue*) and cHi-C contact maps (*red*); (*bottom*) the normalized IS profiles from the models’ (*blue*) and cHi-C contact maps (*red*); (*right*) the distribution of the contact ranks corresponding to the target loops in NPC wt. Notably, the quantities we used to compare models and cHi-C data don’t change considerably in the second half of the simulation indicating that the simulation runs are sufficiently long and numerous to reach a steady state and meaningfully characterize the effect of the parameters on the models’ structural organization.

## Notes

### Competing Interest Statement

The authors have declared no competing interest.

